# An ATF4-centric regulatory network is required for the assembly and function of the OXPHOS system

**DOI:** 10.1101/2023.09.21.558853

**Authors:** Umut Cagin, Manuel José Gómez, Ester Casajús-Pelegay, Rocío Nieto-Arellano, Daniel Arias-Sanroman, Nieves Movilla, Raquel Moreno-Loshuertos, M. Esther Gallardo, Fátima Sánchez-Cabo, José Antonio Enriquez

## Abstract

Identifying the factors that determine mammalian cell viability when oxidative phosphorylation (OXPHOS) function is impaired poses challenges due to the diverse cellular responses and limited clinical material availability. Moreover, animal models often fail to replicate human phenotypes. To address these challenges, this study conducted comprehensive analyses involving multiple defects and species by comparing the RNA-Seq expression profiles of human and murine cell lines with distinct nuclear backgrounds, representing both normal and OXPHOS-deficient models. To minimize species-specific variation, the study employed clustering techniques to group murine genes affected by OXPHOS dysfunction and identified crucial regulators like ATF4, UCP1, and SYVN1. ATF4 consistently displayed activation in response to OXPHOS defects, not only in murine but also in human cells, confirming its pivotal role in the cellular response to mitochondrial dysfunction. By integrating human and murine data, the study unveiled a conserved regulatory network encompassing genes related to the mTOR pathway and folate metabolism. Remarkably, the study uncovered an unexpected finding: the depletion of ATF4 in both mouse and human cells impairs OXPHOS assembly and supercomplex organization. This impairment primarily stems from a severe disruption in complex I assembly in the absence of ATF4, even under non-stress conditions.

## INTRODUCTION

Mitochondria play a central physiological role as cellular metabolic information hub. They collect information about the metabolic status of the cell and their own functional status and receive input from different organelles informing on aspects such as nutrient availability, quality of the protein and nucleic acid metabolism, and disposal of metabolic byproducts. This information is computed to adapt mitochondrial physiology to cellular requirements. A central component of the adaptability of mitochondria is the fact that they have their own genome, their own expression machinery (transcription and translation) and extremely efficient quality control mechanisms devoted to adjust the oxidative phosphorylation system (OXPHOS) to the very different physiological demands that cell differentiation, growth, activity, or death may have under a continuously changing environment ^1^. Integrated expression of the two genomes (nuclear and mitochondrial) that encode components of OXPHOS is important for the effective functioning and plasticity of the bioenergetic system and for many other cellular processes that are integrated with it (metabolism, epigenetics, etc). Mitochondrial DNA (mtDNA) encodes only 13 proteins while the remaining OXPHOS structural components (up to 70) are nuclear encoded, together with the approximately 1,300 characterized proteins located in mitochondria^2^. Physiological or dysfunctional variations in OXPHOS capacity, on the other hand, elicit a nuclear response, known as *Retrograde Response* (RR), that may allow, in the extreme, the survival in culture of cells lacking mtDNA (ρ° cells). The molecular mechanisms responsible for the retrograde response have eluded its determination while more and more evidence connected the failure in the OXPHOS system with the Integrated Stress Response (ISR)^3,4^.

Clinical materials obtained from mitochondrial disease patients, together with model organisms such as mouse, fruit flies and *C. elegans*, have been critical resources to investigate cellular adaptation to OXPHOS deficiency. Mitochondrial communication with the nucleus is the key mechanistic adaptation when the cells’ bioenergetics status is affected. Mito-nuclear communication involves regulation of many conserved signaling pathways including PGC1α, mTOR, Myc, Akt, or Hif1α^5^. For instance, regulation of Hif1α pathway has been shown to ameliorate effects of mitochondrial dysfunction in *Drosophila* motoneurons and can enhance lifespan^4,6^. A valuable work by Khan and colleagues highlighted the importance of performing integrative multi-omics approaches to discover mechanisms and develop therapies for mitochondrial diseases ^7^.

Despite those proposals, an integrated picture of the molecular mechanisms that monitor the status of the OXPHOS system remains to be established. It was expected that a detailed analysis of transcriptomic changes in OXPHOS deficient and OXPHOS rescued cellular models would allow to elucidate the key genes and signaling molecules involved in such monitoring. Transcriptomic technologies have been available already for a while and several groups have applied them to cellular and organismal models of mitochondrial diseases^8–11^. However, those analysis could not provide a complete picture, probably due to several reasons: (i) Most of the earlier studies investigating transcriptomic reprograming upon mitochondrial dysfunction on cellular and organismal models were performed with microarray technologies^6,8–11^; (ii) it has been demonstrated that nuclear background plays a major role in shaping the nuclear transcriptomic reprogramming in the skeletal muscles of mitochondrial encephalomyopathy patients^12^; and (iii) cell transcriptomes are cell type and species specific and the magnitude of the endogenous differences between them overlap with and confuse the interpretation of the differences induced by changes in OXPHOS performance.

To overcome these difficulties, we have analyzed in parallel the transcriptome of cell lines from different nuclear backgrounds, both in human and mouse OXPHOS deficient cellular models, by RNA sequencing. We confirm the strong influence of nuclear background within the same species and between species in two widely used human cell lines (HeLa & 143B), and two mouse cells lines (L929 and NIH3T3), by comparing wild-type OXPHOS function, driven by a common mtDNA, with cells in which mtDNA has been completely lost (ρ°). Other cellular models in which mtDNA has been mutated (mild, partial complex I –CI-deficiency versus complex IV –CIV-depletion in human, and CI or CIV depletion in mice) were also compared to search for common transcriptomic changes that may represent or shed light on potential adaptive response to mtDNA functional alterations. Additionally, to distinguish between adaptation only due to the affectation of the redox status of enzyme cofactors we compared the transcriptomic profiles of OXPHOS deficient mouse cells expressing AOX, the alternative oxidase of *Emericella nidulans*, alone or in combination with NDI1, an alternative mitochondrial NADH dehydrogenase from *Saccharomyces cerevisiae.* Thus, mitochondrial NADH oxidation and electron flux around CoQ is restored without restoring proton-pumping^13^. Using an integrative bioinformatics analysis of these multi-species and multi-conditions data we were able to identify the regulatory network of transcription factors orchestrating transcriptional changes upon mtDNA depletion being the Activating Transcription Factor 4 (ATF4) at its center. In addition, we show that ATF4 deficient fibroblasts undergo a complete reorganization of OXPHOS, and we demonstrate the involvement of ATF4 in the stability of assembled CI and complex (CV), and its requirement for a mammalian cell to survive in the absence of mtDNA.

## RESULTS

### Multi-species cellular models of mtDNA dysfunction

Gene expression profiling of various human and mouse cell lines with mtDNA defects (Table 1) was used to perform a cross-species comparison of the response to mitochondrial dysfunction, at the transcriptional level.

**Table 1.** Summary of cell lines used in the current study. Nuclear and mitochondrial genetic backgrounds are described, as well as phenotypes. Cell lines have been classified into four categories according to the severity of mitochondrial dysfunction.

| Species | Cell line | Nucleus | mtDNA | Phenotype | Severity | Reference |
| --- | --- | --- | --- | --- | --- | --- |
| MOUSE | L929<sup>Rho0</sup> | L929 | $\rho^0$ | Uridine auxotrophy, no functional OXPHOS, depends on glycolysis, no oxygen consumption in SeaHorse | Extreme | 57 |
| MOUSE | NIH3T3<sup>Rho0</sup> | NIH3T3 | $\rho^0$ | Uridine auxotrophy, no functional OXPHOS, depends on glycolysis, no oxygen consumption in SeaHorse | Extreme | |
| MOUSE | L929<sup>COI</sup> | L929 | Point mutation in Col (Complex IV) | Uridine auxotrophy, fails to assemble CIV. Reduced levels of basal respiration with SeaHorse | Severe |  |
| MOUSE | L929<sup>ND4</sup> | L929 | Point mutation in mtND4, dA10227 (Complex I) | Uridine prototrophy. Fails to assemble CI and CI-containing supercomplexes. Severe decrease in O<sub>2</sub> consumption with SeaHorse | Mild | 17 |
| MOUSE | L929<sup>Rho0</sup>AOX | L929 / AOX | $\rho^0$ | L929<sup>Rho0</sup> expressing AOX | Mild | 13 |
| MOUSE | L929<sup>Rho0</sup>NA | L929 / NA | $\rho^0$ | L929<sup>Rho0</sup> expressing NDI1 and AOX | Mild | 13 |
| MOUSE | L929<sup>COI</sup>AOX | L929 / AOX | Point mutation in Col | Respiration capacity independent of complexes III and IV, due to AOX activity | Mild | 13 |
| MOUSE | L929<sup>Balb/c</sup> | L929 | WT | WT | WT | 57 |
| MOUSE | NIH3T3 | NIH3T3 | WT | WT | WT |  |
| HUMAN | 143B<sup>Rho0</sup> | 143B | $\rho^0$ | Uridine auxotrophy, no functional OXPHOS, depends on glycolysis, no oxygen consumption in SeaHorse | Extreme | 14 |
| HUMAN | HeLa<sup>Rho0</sup> | HeLa | $\rho^0$ | Uridine auxotrophy, no functional OXPHOS, depends on glycolysis, no oxygen consumption in SeaHorse | Extreme | 58 |
| HUMAN | 143B<sup>ND4</sup> | 143B | Frameshift in mtND4, iC10947 (Complex I) | Fails to assemble CI and CI-containing supercomplexes. Severe decrease in O<sub>2</sub> consumption with SeaHorse. | Extreme | 16 |
| HUMAN | 143B<sup>LHON</sup> | 143B | Point mutation in mtND4, G11778A (Complex I) | Allows assembly of CI and all supercomplexes. Mild decrease in O<sub>2</sub> consumption with SeaHorse | Mild |  |
| HUMAN | 143B<sup>613</sup> | 143B | WT | WT | WT | 15 |
| HUMAN | HeLa<sup>613</sup> | HeLa | WT | WT | WT |  |

#### Control and mtDNA-less cells

Cells completely devoid of mtDNA, named rho-zero (ρ°) cells, have been intensely used in mitochondrial research for more than 30 years^14^. Here, we choose two human (143B and HeLa) and two mouse (L929 and NIH-3T3) nuclear backgrounds of ρ° cell lines that were compared between them and with cells that had been repopulated with wild type mtDNA from a human donor (613 mtDNA, with a white mtDNA haplogroup H^15^) or mtDNA derived from Balb/c mouse. We confirmed that all four ρ° cells cannot assemble functional OXPHOS, are dependent on uridine supplementation and dependent on glycolysis for energy production (not shown). As expected, oxygen consumption of ρ° (OCR) tested on Seahorse XF analyzer was marginal (Figure1E-F). All four ρ° repopulated cells (control cells), used for the comparative analyses, recovered normal respiration and assembly of OXPHOS complexes. Furthermore, we also used mouse L929 nucleus ρ° cell lines expressing AOX, alone or in conjunction with NDI1 (referred to as NA cells), which had been previously characterized by our group from the energetic and metabolic point of view^13^. *Mutant mtDNA cells*. Because ρ° cells can be considered extreme cases of OXPHOS deficiency, we included additional cell lines, both human and murine, suffering of partial OXPHOS deficiency due to mutations in mtDNA affecting either CI (143B and L929 nuclei) or CIV (only L929 nucleus) function. Human and mouse cells with complex I deficiency were obtained by repopulation of ρ°-143B cells and ρ°-L929 cells, respectively, with mutant mtDNA. In human, we used mitochondria harboring mild or severe CI defects. For mild we used Leber’s Hereditary Optic Neuropathy (LHON) mutant cells with a homoplasmic G to A mutation at position 11778 (143B^LHON^). This mutation affects *MT-ND4* gene and is the most common mutation, found in 69% LHON patients. For severe CI deficiency we used 143B^ND4^ cells (named C4T in former papers^16^) that have a homoplasmic frameshift mutation, consisting in the insertion of an additional C in a row of six Cs at positions 10947-10952, altering the coding sequence of *MT-ND4* and placing a stop codon approximately 150 bp downstream of the mutation^16^. 143B^LHON^ cells exhibit normal levels of CI activity tested by native in-gel activity (Figure 1A). As expected, 143B^LHON^ cells can assemble all OXPHOS complexes and supercomplexes, while 143B^ND4^ cells show no detectable CI activity in gel because they fail to assemble CI and CI-containing supercomplexes (Figure B). Mouse complex I mutant cells L929^ND4^ (named FMI12 in previous papers^17^) have a deletion of an A at position 10227, affecting *mt-Nd4* gene and can survive in the absence of uridine in culture. These cells have no detectable CI activity by in gel activity (Figure 1C) and were able to assemble complexes II, III and IV and supercomplex III_2_+IV, but not CI (Figure 1D). In fact, L929^ND4^ cells have increased III_2_ and III_2_+IV levels derived from the fact that they would associate with CI in normal circumstances (Figure 1D). L929^COI^ mutant mouse cells harbor a homoplasmic missense C6247T point mutation in the cytochrome oxidase 1 (*mt-CoI*) gene that induces a serine to leucine amino acid change; these cells require uridine to survive in culture. Homoplasmic L929^COI^ cells showed residual CIV activity in gel that is concentrated in the monomer CIV, and undetectable supercomplex III_2_+IV activity in the respirasome (Figure 1D). In agreement with that, homoplasmic L929^COI^ mouse cells assemble a very small amount of CIV. Interestingly, the free CIV is strongly reduced and migrates slightly faster in gel than CIV in wild type cells. CIV dimers and I+III_2_+IV supercomplex are dramatically reduced while supercomplex III_2_+IV was present at almost normal levels suggesting that under reduced CIV availability the III_2_+IV supercomplex is preferentially preserved (Figure1D). Interestingly, xenoexpression of AOX in L929^COI^ cells (L929^COI^AOX) induces the elimination of the residual amount of CIV (Figure 1D & see below).

**Figure 1.**
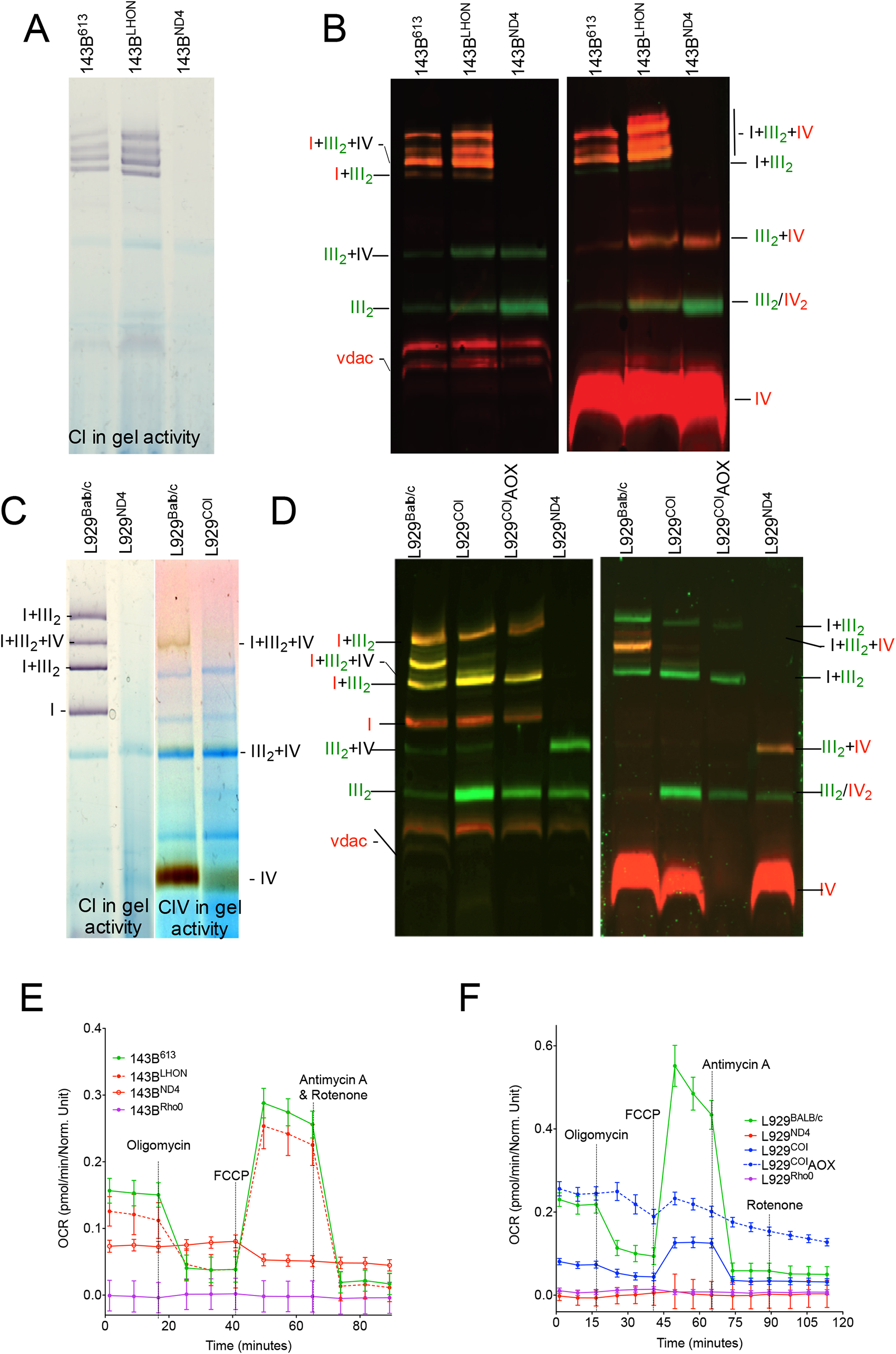
Seahorse and native gel characterization of the indicated human and mouse cell lines. **(A)** Complex I in-gel activity analysis and **(B)** immunodetection of the indicated respiratory complexes and supercomplexes after separation by BNGE of digitonin-permeabilized mitochondria from human 143B-derived cell lines. **(C)** Complexes I and IV in-gel activity assays and **(D)** western blot analysis of the indicated complexes and supercomplexes after BNGE of mitochondria from mouse L929-derived cell lines. **(E-F)** Oxygen consumption rate (OCR) measurement upon sequential addition of oligomycin, FCCP, and rotenone + antimycin A in the indicated human (E) and mouse (F) cells lines using a Seahorse XF96 Extracellular Flux Analyzer.

To obtain a quantitative estimation of the respiration capacity of the different cell lines utilized in this study, we performed respirometry analysis with a Seahorse XF Analyzer. 143B^LHON^ and 143B^ND4^ showed mild decrease and major deficiency in oxygen consumption, respectively (Figure 1E). As expected, homoplasmic L929^ND4^ shows virtually no respiration, while L929^COI^ mutant cells shows a reduced level of basal respiration, very limited spare respiratory capacity and maximal respiration (Figure 1F). Interestingly L929^COI^AOX showed a respiration capacity independent of complexes III and IV due to the additional activity of AOX (Figure 1F). Based on OCR profiles, patterns of mitochondrial supercomplex formation and metabolic properties (uridine dependency) we classified cell lines as having WT, mild, severe, or extreme OXPHOS deficiency phenotype (Table 1).

### Mitochondrial dysfunction severity correlates with transcriptome changes in cell lines with different nuclear backgrounds

It is well documented that the nuclear background of cell lines (depending on species and/or tissue origin) is a major contributor of transcriptional differences. This is also true for mtDNA background *in-vivo*^18^ although no systematic information is available for cultured cell lines. Moreover, very limited information is available on the combined interaction of the three major sources of transcriptome variability. In addition, it is unknown up to which degree the transcriptomic adaptation to mitochondrial dysfunction may be cell type and/or species-specific making it difficult to identify universal responses.

To assess the relevance of nuclear backgrounds in mitochondrial defective human and mouse cells we first estimated the transcriptome differences across different nucleus by RNA Sequencing. Principal component analysis of mouse transcriptomic profiles separated cell lines according to mitochondrial dysfunction severity (Figure 2A). The first dimension (accounting for 35.72% of variability) separated wild type cells (L929^BalbC^ and NIH3T3) and L929^ND4^ (carrying a mutation in gene *mt-Nd4* that results in the loss of CI but maintains uridine-prototrophy) from cells with more severe mitochondrial defects; these were either lacking mitochondrial DNA (ρ° cell lines), or carrying a mutation in *mt-Co1*, independently of whether alternative oxidase (AOX) or NDI1 had been introduced to ameliorate the mitochondrial defect. Analysis of human transcriptomic profiles rendered a similar landscape (Figure 2B). Separation of cell lines according to phenotype severity occurred along the second Principal Component Analysis (PCA) dimension (accounting for 26.74% of variability) because the lack of mtDNA in HeLa background causes a more dramatic adjustment of gene expression programs which was recapitulated by the first dimension (50.35% of variability), which separated ρ° HeLa cells from the other cell lines.

**Figure 2.**
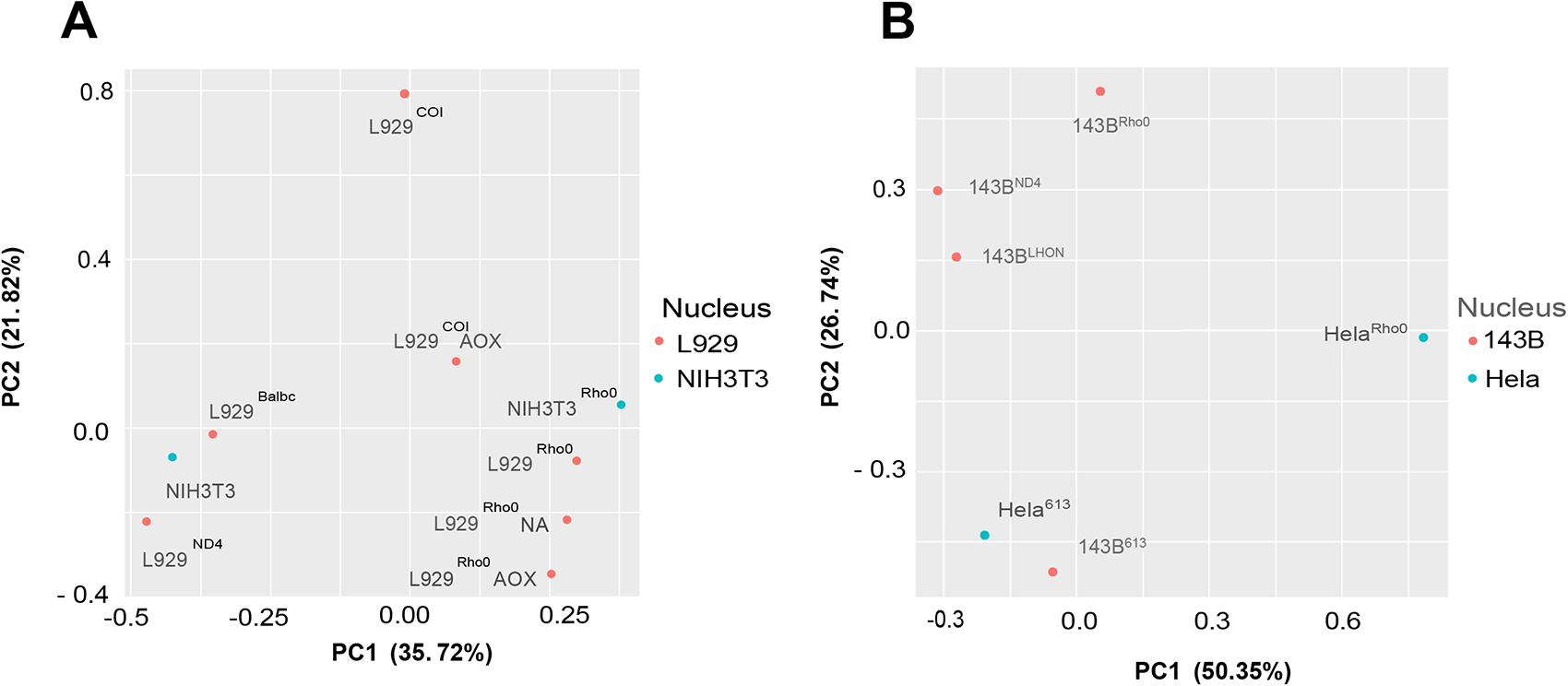
Principal Component Analysis (PCA) of transcriptomic profiles for human and mouse cell lines. **(A)** PCA of mouse transcriptomic profiles separated cell lines according to mitochondrial dysfunction severity. The first dimension (accounting for 35.72% of variability) separated wild type cells (L929-BalbC and NIH3T3) and L929-ND4 (carrying a mutation in gene *mt-Nd4* that results in a mild phenotype) from cells with more severe mitochondrial defects; these were either lacking mitochondrial DNA (ρ° cell lines), or carrying a mutation in cytochrome oxidase I gene (*mt-Co1*), independently of whether alternative oxidase (AOX) or ND1l (an alternative mitochondrial NADH DH from yeast) had been introduced to ameliorate the mitochondrial defect. **(B)** Analysis of human transcriptomic profiles rendered a slightly different landscape. Now, separation of cell lines according to phenotype severity occurred along the second PCA dimension (accounting for 26.74% of variability). The first dimension (50.35% of variability) separate ρ° HeLa cells from the other cell lines, suggesting that the lack of mitochondrial DNA in HeLa background causes a more dramatic adjustment of gene expression programs.

### Unsupervised clustering analysis identifies groups of genes specifically deregulated in mouse cell lines with severe OXPHOS dysfunction

8,008 mouse genes were identified as differentially expressed between control and mitochondrially dysfunctional cell lines (Table S1), confirming a large impact of OXPHOS dysfunction on the transcriptional profile of the cell lines. Unsupervised k-means clustering (k=30) revealed different transcriptional profiles upon different perturbations (Figure S1). *<u>Clusters 1</u>* and *<u>7</u>* contained genes that were either up or downregulated in ρ° cells, respectively. *<u>Cluster 1</u>* was mainly enriched in genes related with one-carbon metabolism, autophagy, and glucose homeostasis, while *<u>cluster 7</u>* genes pointed to oxidative phosphorylation, lipid metabolism and NADH metabolism (Figure S2).

We also identified groups of co-expressed genes that were either up-regulated (clusters 19 and 26) or down-regulated (clusters 13 and 15) in cell lines with severe OXPHOS dysfunction and uridine dependency. Up-regulated genes in *<u>cluster 19</u>* were involved in transport and oxidation-reduction but also in ER response to stress, unfolded protein response, response to hypoxia, autophagy, and metabolism. *<u>Cluster 26</u>* was enriched in genes involved in metabolic processes including glycolysis, gluconeogenesis, carbon metabolism and response to hypoxia. On the other hand, a large proportion of genes down-regulated in OXPHOS-defective cells, *<u>clusters 13</u>* and *<u>15</u>*, were involved in mitosis, DNA replication and repair, nucleotide metabolism and development (Figure S2). Finally, clusters 29 and 10 contained genes that were either up- or down-regulated in cells with any type of OXPHOS deficiency, including L929^ND4^ cells, which had a mild phenotype. For example, genes in *<u>cluster 29</u>* were mostly associated with carbon metabolism and transport, while genes in *<u>cluster 10</u>* were mostly related with lipid metabolism and cell migration (Figure S2).

All in all, our functional characterization of genes clusters suggest that genes deregulated in the absence of mtDNA or defective OXPHOS activate compensatory metabolic and homeostatic mechanisms for the cell to cope with partial or total mitochondrial dysfunction. These processes may involve ER stress, amino-acid metabolism, one-carbon metabolism, and other related pathways.

### ATF4 is the master regulator of metabolic remodeling upon mtDNA dysfunction

With the aim of identifying key regulators responsible for the transcriptional changes observed we determined the most likely upstream regulators for each cluster. We found that the activating transcription factor 4 (ATF4), involved in integrated stress response (ISR), the E3 ubiquitin-protein ligase Synoviolin 1 (SYVN1) and the uncoupling protein 1 (UCP1) were upstream regulators of a significantly enriched set of genes specifically upregulated in ρ° state or in OXPHOS deficient cell lines (clusters 1, 19, 26 and 29) (Figure 3 and Figure S3). Interestingly, other regulators such as the regulator of the ISR Tribbles Pseudokinase 3 (TRIB3) and the transcription factors CREB1 and DDIT3, whose functions are highly related to ATF4^22^, were enriched in at least 2 or 3 clusters (Figure 3). In agreement to this, tunicamycin (a known activator of ATF4^19^) was enriched in clusters 1, 19 and 26 (Figure 3). ATF4 was also identified as a key upstream regulator when predictions were based on the complete collections of differentially expressed genes associated to each relevant pairwise comparison (Figure 4 and Table S2)

**Figure 3.**
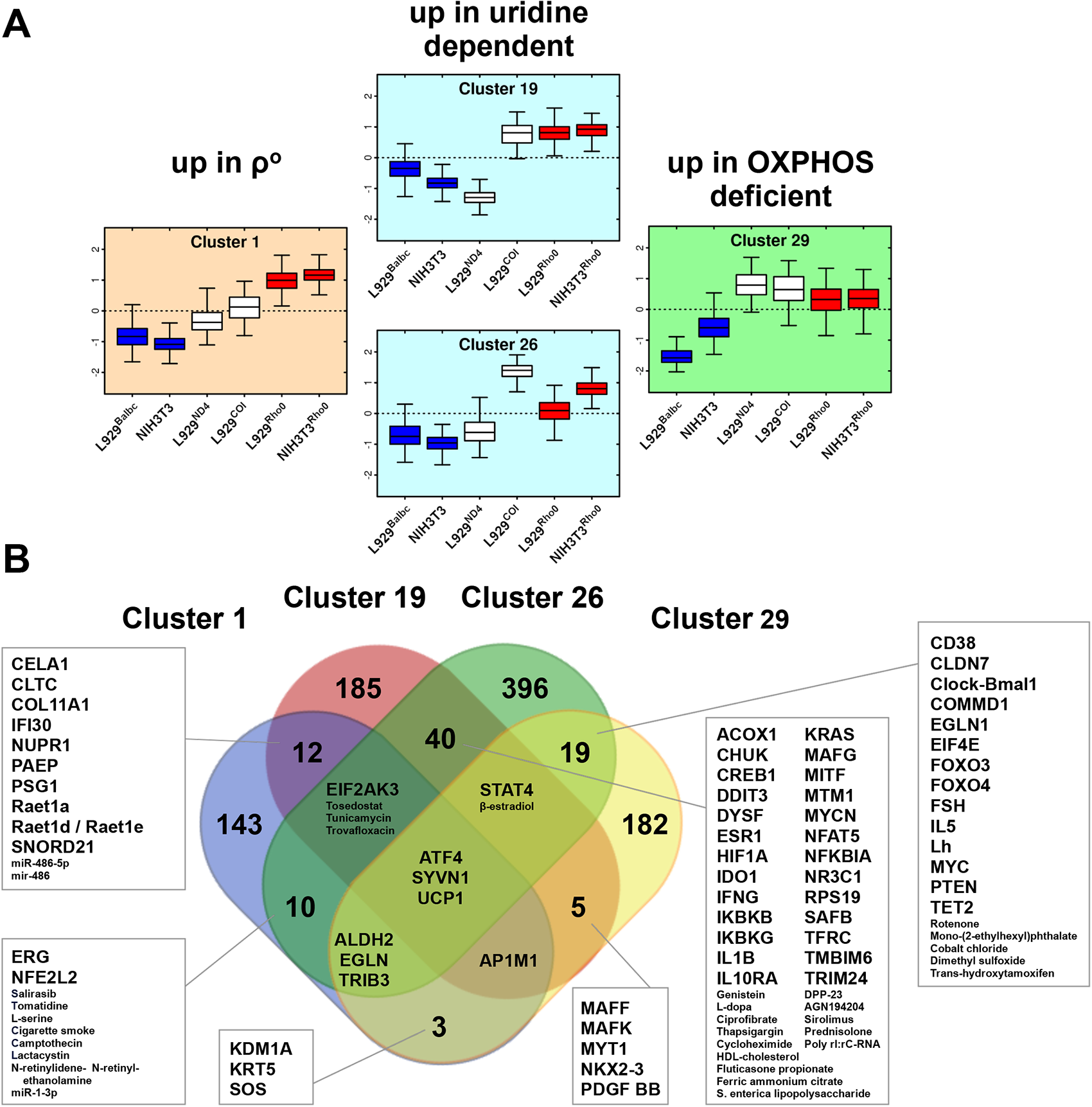
**(A)** collection of 8,008 mouse genes, detected as differentially expressed in contrasts involving control and mitochondrially dysfunctional cell lines, were clustered using k-means, as represented in Figure S1. **(A)** Expression profiles for clusters 1, 19, 26 and 29, which contain genes that are specifically upregulated in ρ° state or in OXPHOS deficient cell lines. **(B)** Summary of IPA-Upstream Regulator analysis results obtained for each of the clusters. ATF4 is the only transcriptional regulator whose targets are enriched in the four clusters.

**Figure 4.**
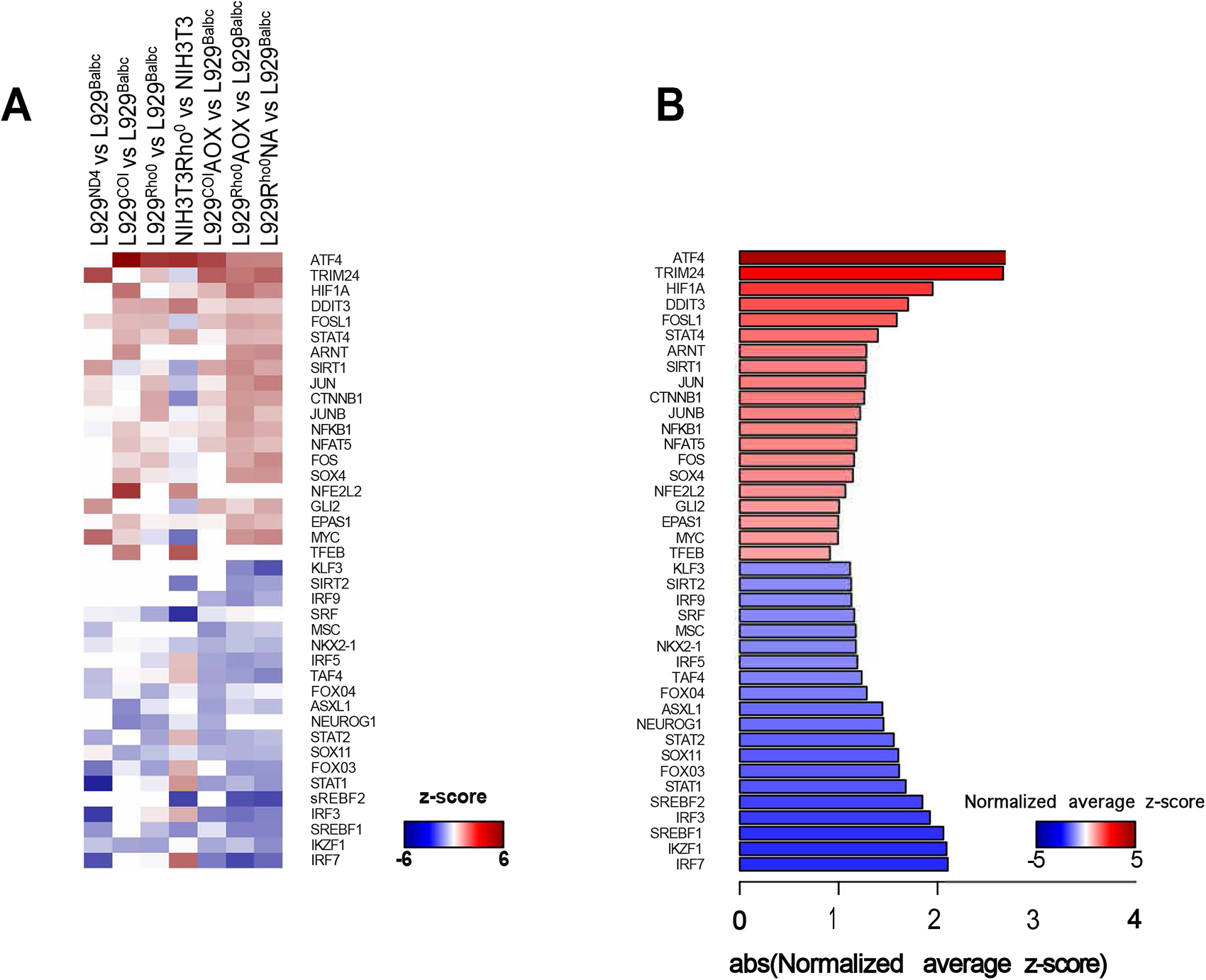
**(A)** Comparative heatmap presenting z-score values for significantly enriched transcriptional regulators, as detected with IPA-Upstream Regulator analysis, on the collections of genes detected as differentially expressed in seven contrasts that compared the expression profile of OXPHOS-deficient mouse cell lines against their control counterparts. **(B)** Average normalized z-score for the same regulators described in panel (A).

To further explore the hierarchy of the regulators involved in the response to mitochondrial disfunction changes, RNA-Seq was carried out to identify differences in the gene expression profile of an ATF4^KO^ mouse embryonic fibroblasts (MEFs) cell line, in the conditions described in Methods. 6,952 genes were identified as differentially expressed, with Benjamini-Hochberg adjusted p-value < 0.05 (Table S3), out of which 3,259 and 3,693 genes were up- or down-regulated, respectively, in ATF4^KO^ cells relative to WT controls. Functional enrichment analyses with IPA and GSEA indicated that differentially expressed genes were mostly associated to developmental functions (Table S3). Twenty-one of the 40 topmost significant upstream regulators identified in pairwise comparisons were expressed in the ATF4^KO^ experiment. Among them, 14 transcription factors (66.67%) were found to be differentially expressed (Table S3) and hence potentially regulated by ATF4. These were SRY-Box Transcription Factor 4 (Sox4) (logFC=14.15), DNA Damage Inducible Transcript 3 (Ddit3) (-2.88), GLI Family Zinc Finger 2 (Gli2) (4.83), Tripartite Motif Containing 24 (Trim24) (2.39), Hypoxia Inducible Factor 1 Subunit Alpha (Hif1α) (-2.41), Endothelial PAS Domain Protein 1 (Epas1) (4.1), Kruppel Like Factor 3 (Klf3) (1.65), Forkhead Box O4 (Foxo4) (1.42), NFKB Inhibitor Alpha (Nfkbia) (1.37), Sirtuin 1 (Sirt1) (1.12), Sterol Regulatory Element Binding Transcription Factors 1 (Srebf1) (-0.97) and 2 (Srebf2) (0.70), JunB Proto-Oncogene, AP-1 Transcription Factor Subunit (Junb) (0.68). In fact, the network representation of the differentially expressed targets of each TF unbiasedly identified Atf4, Hif1a, Ddit3, Sirt1 and NfkbIA as the network hubs with the highest closeness (circle size), what indicates their influential position in the network (Figure 5).

**Figure 5.**
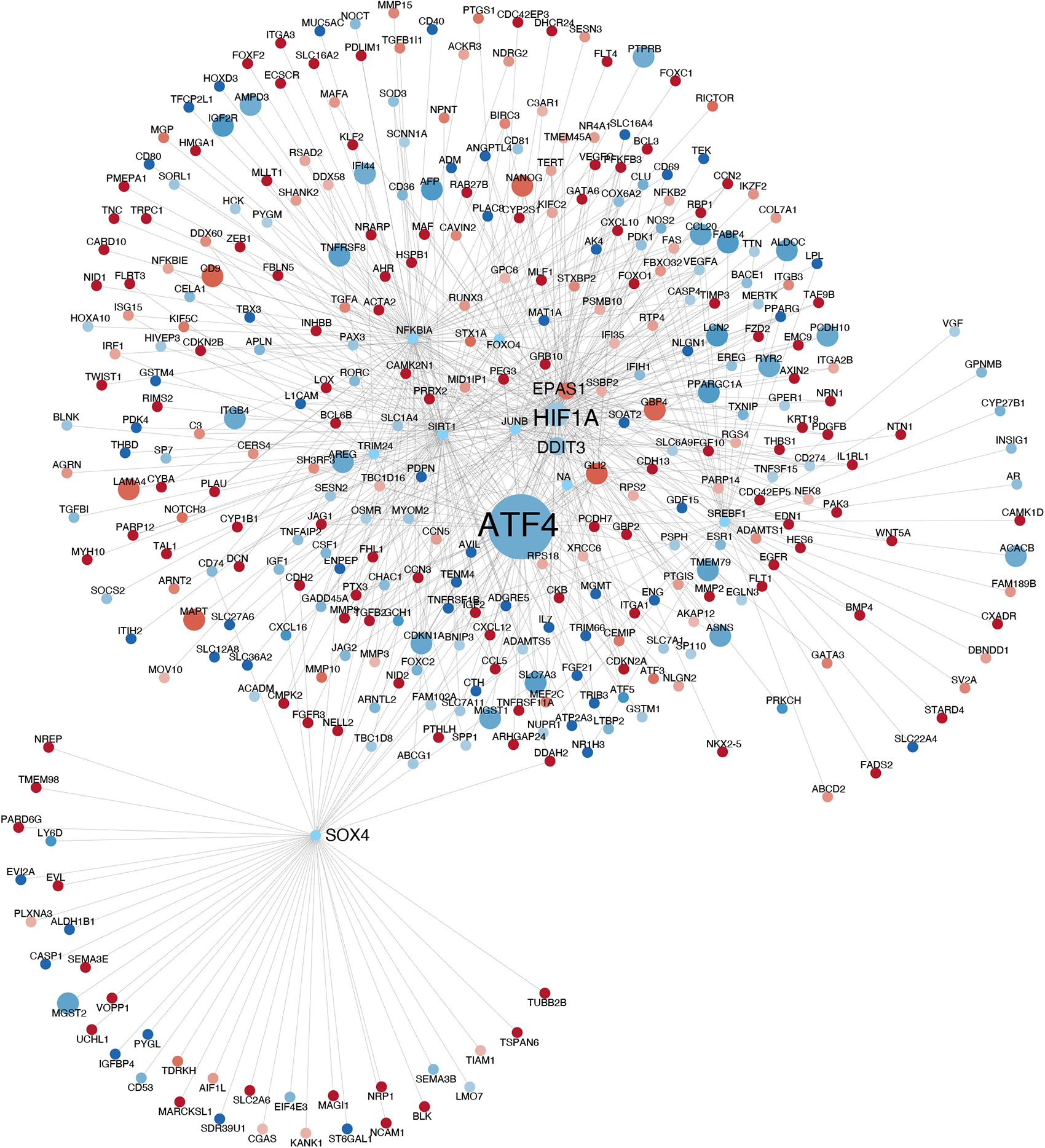
Gene network summarizing the relations between enriched transcriptional regulators and its targets, as derived from IPA-Upstream Regulator analysis results for of 6,952 differentially expressed genes detected in the comparison of ATF4^KO^ mouse embryonic fibroblasts (MEFs) cell line and their control counterparts. Nodes are colored by logFC values, and their size represents the betweenness of each node.

On a different side, 48 OXPHOS genes, out of the collection of 168 genes defined in MitoCarta 3.0, were differentially expressed in the ATF4^KO^ cell line, relative to controls (Fig S4A). To explore additional connections between the Atf4 network and OXPHOS genes, we predicted binding sites for the transcription regulators included in the network. TF binding sites were detected in the promoters of 48 out 168 OXPHOS genes, and in 14 of the 48 OXPHOS genes that had been detected as differentially expressed in the ATF4^KO^ versus control comparison (Fig S4 B).

### Activation of ATF4 is required for survival upon mitochondrial stress

Cell lines used in the analysis were stable cell lines and already adapted to culture conditions. For example, mtDNA depleted (ρ°) cells were obtained by long-term treatment with ethidium bromide (EtBr). To check the dynamic nature of the Atf4 response to mtDNA dysfunction, control (L929^Balbc^) cells were treated with EtBr for 72 hours to obtain partial depletion of mtDNA. After 72 hours, ATF4 protein levels were dramatically increased (Figure S5A). Treatment with rotenone, a known inhibitor of CI, also resulted in increased levels of ATF4 (Figure S5A), in agreement with a study using oligodendroglia^20^. We also tested whether ATF4 expression is affected upon defects in mtDNA translation. We found that treatment with doxycycline, an inhibitor of mtDNA translation^21^, results in increased ATF4 levels in mice control L929^Balbc^ cells (Figure S5A). Interestingly, ATF4 levels were correlated with doxycycline treatment in a time-dependent manner. Similar results have been obtained in a study using HeLa cells^22^.

We then used ATF4^KO^ MEFs to obtain information regarding its sensitivity to the same collection of drugs. ATF4^KO^ mouse cells were more sensitive to EtBr, rotenone and doxycycline treatment compared to control L929^Balbc^ cells (Figure S5B). Of note, we failed to obtain ρ° derivatives from the ATF4^KO^ cell line, since long term treatment with EtBr resulted in the death of all cells in 4/5 months. This further proves that ATF4 upregulation and activation is necessary for the survival of cells in the absence of mtDNA or defective OXPHOS.

### Cross-species analysis reveals an ATF4 centric transcriptional network involved in a conserved response upon mitochondrial defects

Following the analysis in mouse cell lines, unsupervised clustering was performed on a collection of 8,165 DEGs identified after transcriptome profiling of human cell lines (Table S1). As before, we used k-means to define 30 clusters of co-expressed genes, which were subject to Upstream Regulator analyses with IPA. Interestingly, ATF4 was enriched in four clusters (Cluster 16, 23, 28 and 30) (Figure S6) indicating the prominent role of ATF4 in metabolic adaptation also in human cells. More importantly, the shapes of those clusters, representing patterns of gene expression differences associated to various states of mitochondrial dysfunction, were highly like gene clusters observed in mouse, indicating that a similar gene regulatory network may exist in both species.

To search for a conserved shared network where ATF4 could be involved, we generated two meta-clusters, MCOMB and HCOMB, which combined murine and human clusters, respectively, that had been identified as enriched in ATF4 targets after applying a stringent significance threshold (adjusted p-value < 0.05) (Figure 6A). After performing Upstream Regulator analysis on both meta-clusters, we identified eight transcription factors: Atf4, Ddit3, Hif1α, Hepatocyte Nuclear Factor 4 Alpha (Hnf4α), Nfkbia, Nuclear protein 1 (Nurp1), Signal Transducer and Activator of Transcription 4 (Stat4) and Tumor Protein P53 (Tp53) whose targets had been detected as highly enriched in both meta-clusters (Figure 6B). Atf4 was the transcriptional regulator enriched with the maximal significance in both gene sets (Figure 6C). This was expected, of course, given that individual clusters had been selected because they were enriched in ATF4 targets. However, the fact that mouse and human meta-clusters were enriched in targets of an additional common set of seven regulators was very relevant because it suggested the existence of a common regulatory network (referred here as the Atf4 network) involved in a conserved response to mitochondrial OXPHOS dysfunction. To summarize the functions in which the set of shared regulators could be involved, we defined eight collections of genes, representing a total of 1,325 orthologous genes that had been detected as differentially expressed both in human and mouse and that, at the same time, were targets of some of the eight shared regulators. Each collection of genes was subjected to enrichment analysis to identify functional terms that are connected to each of the regulators (Figure 6D). Results indicated that the set of eight transcriptional regulators is not only involved on mitochondrial function regulation, but also on the regulation of ER-stress, calcium homeostasis and several aspects of energy metabolism. In summary, these results together show that ATF4 activation is a conserved signal in response to mitochondrial dysfunction both in human and mouse cells.

**Figure 6.**
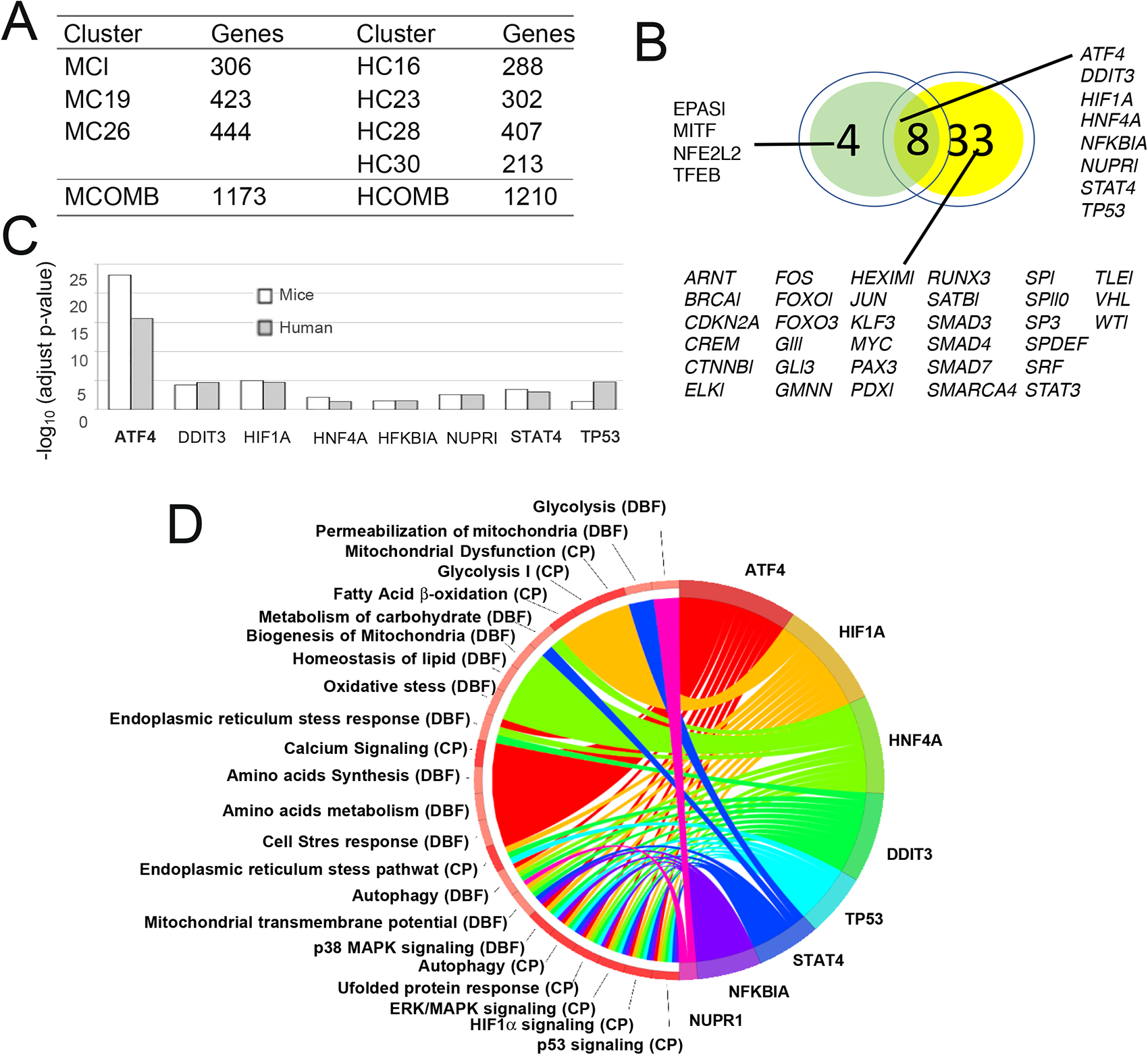
**(A)** Mouse and human clusters enriched in ATF4 targets (with adjusted p-value < 0.05; MC* and HC*, respectively), and combined gene sets in mouse and human (MCOMB and HCOMB, respectively). **(B)** IPA-Upstream Regulator analysis results summary for MCOMB (green) and HCOMB (yellow), with indication of a set of eight shared transcriptional regulators. **(C)** Enrichment significance for the eight shared transcriptional regulators in human and mouse. **(D)** Enriched selected functions from the Canonical Pathways (CP) and Diseases and Biofunctions (DBF) IPA ontologies, associated to the target genes of each of the eight shared regulators

### ATF4 deficient cells have defective OXPHOS assembly and performance

ATF4 is required for Integrated Stress Response (ISR) and more specifically its upregulation is a classical marker for ER-stress^23^. Therefore, ATF4 defective cells cannot adapt to stress conditions affecting ER homeostasis. It has been recently shown that ATF4 is also required for expression of UPR genes and that loss of ATF4 results in enhanced oxidative damage and reduced mitochondrial membrane potential^24^. ATF4^KO^ MEFs showed normal growth pattern in DMEM supplied with essential and non-essential amino acids, and 50 µm β-mercaptoethanol. However, they could not survive in media without these supplements. ATF4^KO^ MEFs had reduced levels of basal and maximal respiration compared to L929^Balbc^ control cells and their spare respiratory capacity was very low (Figure 7A). Blue Native-Polyacrylamide Gel Electrophoresis (BN-PAGE) analysis showed a re-organization of the whole OXPHOS system (Figure 7B). More specifically, reduced levels of CI and CI-containing supercomplexes (I+III_2_ and I+III_2_+IV) and slightly less CIV and CV were detected. The highest MW band, which may represent a dimmer of I+III_2_, is completely absent in ATF4^KO^ cells. On the contrary, due to the re-organization of the OXPHOS system, more III_2_ and III_2_+IV supercomplexes were detected. Interestingly, ATF4^KO^ cells accumulate low molecular weight CV subunits which may represent a partially assembled CV (Figure 7B). The OXPHOS system re-organization phenotype is reminiscent of CI OXPHOS deficiency. Therefore, we analyzed native CI in-gel activity and detected reduced levels of CI activity (Figure 7B).

**Figure 7.**
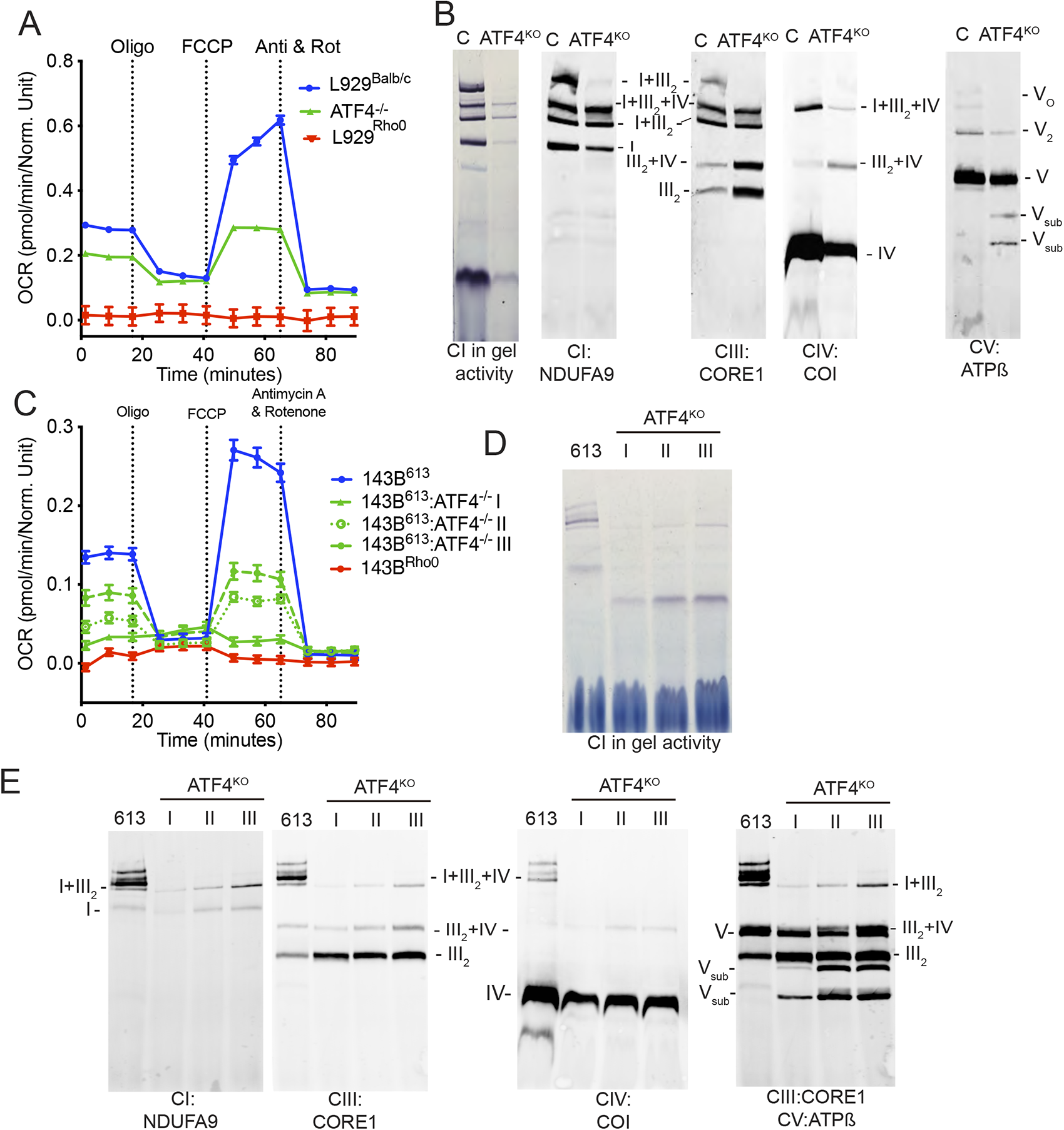
Seahorse and native gel characterization of mouse ATF4^KO^ MEFS and human ATF4^KO^. **(A-B)** Oxygen consumption rate measurement in mouse ATF4^KO^ cells compared to L929^Balb/c^ and L929^Rho0^ cell lines **(A)** and OXPHOS complexes and supercomplexes organization analysis in mouse ATF4KO cells compared to L929Balb/c **(B)**. **(C-E)** Respiration activity of three independent clones of human 143B derived ATF4^KO^ cells measured using the Seahorse technology **(C)** and pattern of mitochondrial supercomplexes analyzed by BNGE followed by complex I in-gel activity **(D)** and immunodetection of the indicated respiratory complexes subunits **(E)**

To generalize the role of ATF4 in the regulation of CI, we then generated ATF4^KO^ human cell lines from 143B^613^ control cells by Crispr/Cas9 gene editing. After targeted nuclease expression and single cell sorting of GFP-positive cells, we characterized several single-cell colonies (named Clone I, II and III) for which the absence of the last part of ATF4 exon 1 was proven by PCR (not-shown). As expected, these colonies could not survive in the absence of β-ME (Figure S7A) and had undetectable levels of ATF4 protein even after treatment with doxycycline (Figure S7B). These three ATF4^KO^ clones had reduced oxygen consumption rates (Figure 7C). Complex I in-gel activity assay reflected strong reduction in CI activity, correlating with the oxygen-consumption rates (Figure 7D). Human ATF4^KO^ cells reproduced the reduction in CI amount and the accumulation of low molecular weight-CV subassemblies observed before in ATF4^KO^ mouse cells (Figure 7E). In summary, ATF4 is not only required to cellular adaptation to OXPHOS defect but also for the normal biogenesis and functionally competent OXPHOS systems both in human and mouse cells.

### ATF4 controls circadian rhythm

The requirement of ATF4 for the correct biogenesis of the OXPHOS system was somehow surprising since this transcription factor is barely detected in wild type cultured cells under non-stress conditions. We hypothesized that the relevance of this transcription factor may be related with the requirement of ATF4 to the proper expression of circadian master regulatory genes and the fact that its expression follows also a period^25,26^. Moreover, mounting evidence suggest that the biogenesis of the OXPHOS system and its recycling is also circadian^27–29^. In agreement with this proposal, we identified 154 expressed putative circadian related genes in our RNA-Seq, out of 229 mouse genes annotated as “circadian” in the Gene Ontology database. Thus, 83 of them (53.8%) were differentially expressed between WT and ATF4^KO^ cells, including Clock and Bmal1 (Table S3). This observation confirms the requirements of basal ATF4 activity for a proper cellular circadian function.

## DISCUSSION

Heterogeneity of mitochondrial disorders has been a serious drawback in the diagnosis and treatment of these detrimental diseases. Limited availability of clinical material has also limited the amount of genetic information that can be obtained from patients. Information scarcity is revealed by the fact that many published datasets are only partially overlapping, if not contradictory, a constraint that could be sorted by performing multi-defect and multi-species comparative analyses. On a different side, and from a methodological point of view, advances in sequencing technologies, especially exome sequencing, helped clinicians to characterize disease causing mutations in mitochondrial related genes^30^. In addition, transcriptomic technologies, especially RNA Sequencing, have served as a very important tool to uncover the roles of such genes. Most of the studies investigating transcriptomic changes associated to mitochondrial dysfunction, however, have been carried out with microarray technologies. In fact, the only published systematic approach incorporated datasets from several organisms (*H. sapiens*, *M. musculus*, *D. melanogaster* and *C. elegans*) and various microarray platforms^31^.

In the current study we have compared the expression profiles of human and murine cell lines with different nuclear backgrounds, representing WT or OXPHOS deficient cellular models. To eliminate some sources of variability, and to be in position of reaching more conclusive and trustable conclusions, we have worked exclusively with transcriptomic datasets generated ‘in-house’, by RNA-Seq. As it could be expected, we have observed significant differences at transcriptomic level in the response to OXPHOS dysfunction in human and mouse cell lines. Nuclear background specific changes have also been observed, especially in human cells. One should consider that mouse cell nucleus derives from close related and inbred strains that have lost heterozygosity and variability. The human, however, are derived from different individuals and present heterozygosity and genetic variability. In our case, for example, HeLa cell nucleus derive from an Afro-American female while the 143B nucleus comes from a white individual. This in turn may explains the limited consensus on the published datasets.

To reduce the species-specific transcriptomic variation, we initially focused on murine cell lines having defective mitochondria. K-means clustering of murine genes whose regulation was altered by OXPHOS dysfunction was used to identify groups of genes whose expression level correlated with the severity of OXPHOS defects. Downstream functional analyses on these clusters allowed us to identify three regulators: ATF4, UCP1 and SYVN1 (Figure 3). ATF4 is the only transcription regulator in this list since UCP1 is a carrier and SYVN1 is a ligase. Furthermore, ATF4 was also detected as a highly activated and enriched transcription factor in most pairwise comparisons between OXPHOS deficient cell lines and their corresponding WT counterparts (Figure 4A). Interestingly, enrichment of ATF4 targets was also detected in the collections of dysregulated genes from cell lines expressing AOX or NdiI/AOX. In these cells the rescue of the mitochondrial function was not fully achieved by the expression of AOX and NdiI (no recovery of proton pumping). Therefore, ATF4 activation is involved in the response to OXPHOS defects, independently of their severity. Supporting this, we have detected upregulation of ATF4 at the protein level upon chemically inducing mitochondrial dysfunction *in-vitro* by EtBr, rotenone, and doxycycline treatments.

ATF4’s novel role in regulating one-carbon metabolism upon mitochondrial dysfunction has been recently uncovered by several studies in cellular and animal models^2,4,7,19,32–35^. In correlation with our results on ρ° cell lines, TRIB3 expression level was also among the top 5 highest expressed genes in a microarray-based study that characterized the expression profile of human mtDNA depleted cells^36^. Of note, TRIB3 induction is observed upon variety of stress conditions including oxidative stress, ER stress, glucose and amino acid deficiency^37^.

After using k-means to cluster human genes whose regulation was altered by OXPHOS dysfunction, we were able to identify a set of clusters that were enriched in ATF4 targets. Upstream regulator analysis of murine and human meta-clusters, defined by the combination of ATF4-target enriched clusters in mouse and human, respectively, resulted in the identification of a common set of eight regulators, including ATF4. We propose that these genes conform a regulatory network conserved between mouse and human cells, which is activated upon mitochondrial dysfunction (Figure 5). This network includes regulators such as Hif1a, highlighting the interconnected nature of oxidative stress and mitochondrial dysfunction. Interestingly, this conserved regulatory network had overlapping with the regulatory elements with the Integrated Stress Response (ISR) such as DDIT3, therefore confirming the role of ISR as a key signaling element, conserved between two species and various mitochondrial deficiencies: mtDNA loss, CI and CIV mutants.

It has been shown that ATF4 itself can regulate purine synthesis by interacting with mTORC1^38^. In agreement with this study, we have observed the enrichment of genes related to mTOR pathway in most of the collections of genes that are differentially regulated in association to OXPHOS dysfunction. One-carbon metabolism plays a central role in biosynthesis of purines and thymidine, amino acid homeostasis, redox defense and methylation^39^. Folate-mediated one-carbon cycle is compartmentalized both in cytosol and mitochondria. Since mitochondria determine NAD^+^/NADH ratio within the cell, regulation of one-carbon metabolism can have important energy buffering roles. Specifically, it has been shown that folate metabolism has direct roles in mitochondrial NADPH production via ALDH1L2 (Aldehyde Dehydrogenase 1 Family Member L2, Mitochondrial 10-Formyltetrahydrofolate Dehydrogenase) and MTHFD2 (methylenetetrahydrofolate dehydrogenase 2), therefore can balance the changes in NAD^+^/NADH ratio^40^. Furthermore, we have observed conserved MTHFD2 activation, as shown in the table in Figure 4, supporting the folate metabolism activation as a metabolic survival mechanism^38^. Similar approaches of characterization of metabolic deficiencies that are associated to mitochondrial dysfunction can also uncover new therapeutic avenues, as exemplified by the discovery that enhancing nucleotide metabolism has beneficial effects in *PINK1* mutant animal models^41^. Tissue specific metabolic requirements, especially related to one-carbon metabolism^42^, may be the underlying cause of heterogeneity observed in mitochondrial patients. Although it requires more research, we believe folate (folic acid), or nucleotide supplementation may have beneficial effects on mitochondrial patients with defects in mtDNA (mutation or depletion).

It has been recently shown that ATF4 silenced cells have reduced mitochondrial membrane potential and increased oxidative stress^24^. Although several metabolic roles of ATF4 have been recently discovered, its direct role on OXPHOS assembly was not notice. We observed complete re-organization of OXPHOS and reduced CI levels and activity. These observations are in-line with these cells being more sensitive to rotenone, doxycycline and EtBr treatment. We postulated that the ATF4 dependent CI-biogenesis may be indirectly controlled by the circadian role of ATF4 in normal cells. Our proposal is in agreement with the observation that mitochondrial OXPHOS activity is rhythmic and regulated by CI acetylation under the control of clock genes^43,44^.

Surprisingly, accumulation of low molecular-weight CV bands on BN-PAGE may suggest that ATF4 has wider roles in CV assembly or plasticity. Interestingly, a recent report by Balsa and colleagues showed that ATF4 can regulate OXPHOS supercomplex organization upon endoplasmic reticulum (ER) and nutrient stress via activation of PERK-eIF2α axis^45^. Importantly, the same study reported increased ATF4 levels upon galactose treatment in cultured U2OS cells. In summary, the exact involvement of ATF4 in the assembly of OXPHOS complexes is not fully known yet and requires further investigation.

This work not only provides mechanistic insights into the response to mitochondrial dysfunction but also information that may affect the choice of clinical markers for mitochondrial diseases. Non-invasive methods for mitochondrial patient diagnosis are very critical since muscle biopsies are not an easy procedure for mitochondrial patients with myopathies. Until recently, only ‘serum lactate’ measurements were consulted for many years as a marker for mitochondrial patients. FGF-21 has been demonstrated as a promising biomarker for mtDNA defected myopathies^46^, and it is also regulated by ATF4. More recently, the activation on FGF-21 has been demonstrated in neuronal mitochondrial dysfunction following ablation of the mitochondrial fission protein Drp1^47^. Here, we further prove that ATF4 centered biomarkers studies can be more trustable, putting FGF-21 and serine levels at the center of the stage in understanding and treating mitochondrial diseases.

## Acknowledgments

We acknowledge all GENOXPHOS group members for their scientific discussions contributing to this manuscript. We thank Dr. David Ron for providing ATF4^-/-^ mouse cells. We thank members of the CNIC facilities (Genomics, Bioinformatics, Cell culture, Animal facility and Viral vectors) for their technical assistance.

## Author contributions

JAE & UC conceived and designed experiments. U.C. performed CRISPR KO, respirometry, enzymatic activity, mitochondrial isolation, BN-gel and immunoblot analysis, assisted by RN-A, EC-P and DA-S in different stage of the experimental work. FSC & MJG performed quantitative analyses of RNAseq and computational network modeling experiments. Data analysis and figure creation was carried out by UC, MJG, FSC and J.A.E. that also wrote the manuscript with input from all authors.

## Methods

### Cell Lines and Media

All cell lines were grown in DMEM (D5796 Sigma-Aldrich) supplemented with 5% FBS (fetal bovine serum, GIBCO-BRL) and 1 mM sodium pyruvate (Lonza) in a 5% CO_2_, 95% air atmosphere at 37°C. Cell lines lacking mtDNA (ρ°) and mtDNA mutants were supplemented with 50 μg/ml uridine. Most cell lines used in this study were generated and described previously (Table 1). For drug treatments, medium containing the indicated concentration of Rotenone (Sigma), BFA (Sigma), Doxycycline (Sigma, D9891) were used.

ATF4^-/-^ mouse cells (MEFs) used in this study were described earlier ^48^. ATF4 mutant human osteosarcoma cell lines were obtained from control 613 cells by CRISPR/Cas9 gene editing technology. GuideRNA (gRNA) were selected to drive the elimination of the last portion of exon 1. After each round of Crispr/Cas9 endonuclease expression, single cells expressing GFP were collected by cell sorter and colonies were genotyped by PCR. Sequences of primers used for genotyping of the colonies were: atgatggcttggccagtg (forward) and ccattttctccaacatccaatc (reverse). Two rounds of CRISPR/Cas9 expression were applied to mutate both alleles. A third round of expression and single cell sorting and genotyping was applied to minimize any possibility of having mixed culture population. Furthermore, viability of these colonies was tested by media lacking βME, EAA and NEAA.

### Western Blotting

After electrophoresis (SDS-PAGE, BNGE, or BN-SDS-PAGE), gels were electroblotted onto Hybond-P polyvinylidene fluoride (PVDF) membranes (GE Healthcare) and sequentially probed with specific antibodies against ATF4 (Cell Signaling ATF-4 (D4B8), #11815) and GAPDH (Abcam). Secondary antibodies were peroxidase conjugated (Invitrogen) when the signal was generated using ECL Plus (GE Healthcare) or conjugated to LI-COR IRDye 800CW or IRDye 680LT when the signal was acquired with the ODYSSEY Infrared Imaging System (LI-COR). The relative amount of each band was estimated with GelEval software from the scanned membranes or with the ODYSSEY Infrared Imaging System (LI-COR).

### BN-PAGE and In-gel Activity of OXPHOS Complexes

Mitochondria were isolated, run on 5-13% gradient gels and analyzed by BNGE, according to Wittig et al.^49^, with some modifications^50^. SDS-PAGE was conducted with strips excised from the first BNGE dimension. Antibodies that were used to compare the levels of OXPHOS complexes are: CI-Ndufa9 (Abcam), CII-Fp70 (Molecular Probes), CIII-Core1 (Abcam), CIII-UQCRC2 (Abcam), CIV-CoI (Invitrogen), CV-ATPB (Abcam). VDAC1 (Mitosciences) was used as a loading control antibody. In order to determine CI in-gel activity, NADH dehydrogenase activity was measured in isolated mitochondria. BN gels were incubated in a buffer containing 0.1 M Tris-HCl, 0.14mM NADH and 1mg/ml Nitro Blue Tetrazolium overnight at 25 °C

### Respirometry and Oxygen Consumption with Seahorse XF Analyzer

Oxygen consumption was measured with an XF96 Extracellular Flux Analyzer (SeaHorse Bioscience), as specified by the provider. Mito Stress Kit was used as a standard way of assessing mitochondrial OXPHOS performance. Data were normalized to DNA content with the CyQuant NF Cell Proliferation Assay Kit (Molecular Probes, C35006).

### Cell viability and proliferation

To assess cellular proliferation, growth curves were obtained by seeding 3,000 cells per well in 96-well plate. At least 8 replicate wells were used per condition. Cyquant (Molecular Probes, C35006) was used as a quantifiable measurement for cell number.

### RNA Isolation and RNA Sequencing

Cells were grown on p100 cell culture dishes until reaching approximately 80% confluency. The plate was washed with PBS and cells were collected immediately. Total RNA was extracted with TRIzol reagent and then purified on RNeasy spin columns (Qiagen). Total RNA was quantified, and purity checked using a NanoDrop ND-1000 (Thermo Scientific, Waltham, MA, USA). RNA integrity (RNA Integrity Score ≥ 7.9) and quantity were determined with an Agilent 2100 Bioanalyzer. 500 ng of total RNA were used with the TruSeq RNA Sample Preparation v2 Kit (Illumina, San Diego, CA) to construct index-tagged cDNA libraries. Libraries were quantified using a Quant-iT™ dsDNA HS assay with the Q-bit fluorometer (Life Technologies, Carlsbad, California). Average library size and the size distribution were determined using a DNA 1000 assay in an Agilent 2100 Bioanalyzer. Libraries were normalized to 10nM using Tris-HCl 10mM, pH8.5 with 0.1% Tween 20. Libraries were applied to an Illumina flow cell for cluster generation (True Seq SR Cluster Kit V2 cBot) and sequence-by-synthesis. Single reads (75 base long) were generated using the TruSeq SBS Kit v5, on Illumina platforms Genome Analyzer IIx or HiSeq 2500, following the standard RNA sequencing protocol. Reads were further processed using the CASAVA package (Illumina) to demultiplex reads according to adapter indexes and produce fastq files.

### Bioinformatic Analysis of RNASeq data

Fastq files were pre-processed with a pipeline that used Cutadapt 1.2.1^51^ to remove TruSeq adaptor remains, and FastQC^52^ to perform quality checks on the reads. Then, RSEM^53^ was used to align pre-processed reads against transcriptome references GRCm38.v76 or GRCh38.v76, and to obtain expression estimates at gene level. Raw count matrices were used as input for a differential expression pipeline that used ComBat^54^ to correct for batch effects and limma^55^ for normalization and differential expression testing, taking into consideration only nuclear genes with at least 1 count per million in at least two samples. Genes were classified as differentially expressed if changes were associated to Benjamini-Hochberg adjusted *P*-value < 0.05 and abs(logFC) > 0.5 in any of the pairwise contrasts performed, in which the transcriptomic profile of OXPHOS deficient and WT cell lines were compared (Table S1). Raw reads and TMM-normalized batch corrected counts have been deposited in GEO with the accession number GSE. The gene network was built using the information on the upstream regulators identified for IPA from the set of differentially expressed genes between ATF4^KO^ vs WT mice. Only genes differentially expressed between both conditions were considered in the analysis.

### KMeans Clustering and Downstream Functional Analysis

To identify groups of genes with similar expression profiles across the cell lines being considered, k-means clustering was performed with the R ComplexHeatmap package on sets of 8,008 murine genes or 8,165 genes human genes that had been identified as differentially expressed in a selection of pairwise contrasts (Table S1). The elbow method was used to estimate the optimal number of clusters, which was set as k=30, both for mice and human samples. Metaclusters were generated by manually merging selected clusters. Gene clusters and metaclusters were functionally annotated with DAVID^56^, to identify significant associations to Biological Process GO terms and KEGG pathways, as well as with the Upstream Regulator tool of Ingenuity Pathway Analysis (IPA, Qiagen), to identify potential regulators. IPA was also used to recover functional interactions between the set of eight regulators shared by mouse and human metaclusters, and to recover information about associated functions.

### Statistical Analysis and Graphical Representation

Statistical analyses and graphics were produced with GraphPad Prism 6 software or with the R suite. Data sets were compared by unpaired two-tailed *t*-tests. Differences were considered statistically significant at *P* values below 0.05. \**P* < 0.05; \*\**P* < 0.005; \*\*\**P* < 0.0005. All results are presented as mean ± s.d. or mean ± s.e.m.

## Supplemental Information

### Supplementary Tables

**Table S1.**
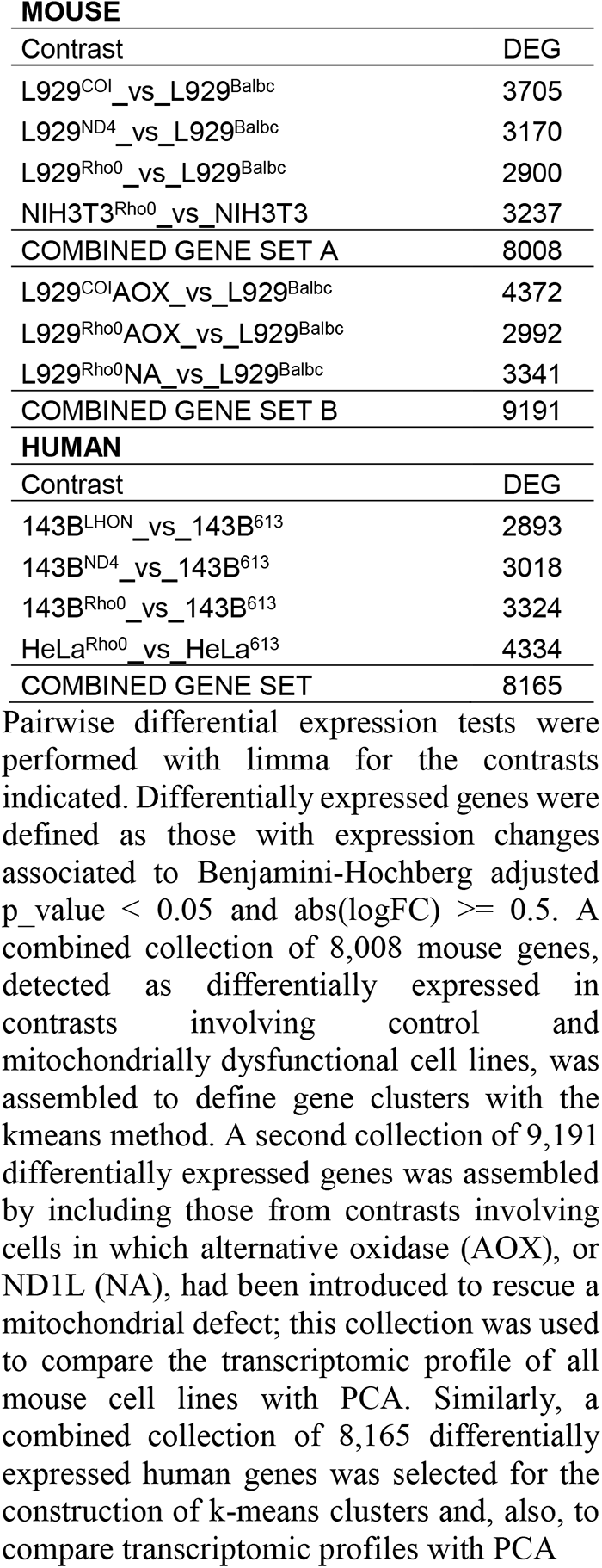
Summary of contrasts and numbers of differentially expressed genes, and combined sets of differentially expressed genes, in human and mouse.

| **MOUSE** |  |
| --- | --- |
| Contrast | DEG |
| L929<sup>COI</sup>_vs_L929<sup>Balbc</sup> | 3705 |
| L929<sup>ND4</sup>_vs_L929<sup>Balbc</sup> | 3170 |
| L929<sup>Rho0</sup>_vs_L929<sup>Balbc</sup> | 2900 |
| NIH3T3<sup>Rho0</sup>_vs_NIH3T3 | 3237 |
| COMBINED GENE SET A | 8008 |
| L929<sup>COI</sup>AOX_vs_L929<sup>Balbc</sup> | 4372 |
| L929<sup>Rho0</sup>AOX_vs_L929<sup>Balbc</sup> | 2992 |
| L929<sup>Rho0</sup>NA_vs_L929<sup>Balbc</sup> | 3341 |
| COMBINED GENE SET B | 9191 |
| **HUMAN** |  |
| Contrast | DEG |
| 143B<sup>LHON</sup>_vs_143B<sup>613</sup> | 2893 |
| 143B<sup>ND4</sup>_vs_143B<sup>613</sup> | 3018 |
| 143B<sup>Rho0</sup>_vs_143B<sup>613</sup> | 3324 |
| HeLa<sup>Rho0</sup>_vs_HeLa<sup>613</sup> | 4334 |
| COMBINED GENE SET | 8165 |

Pairwise differential expression tests were performed with limma for the contrasts indicated. Differentially expressed genes were defined as those with expression changes associated to Benjamini-Hochberg adjusted $p\_value < 0.05$ and $abs(logFC) \geq 0.5$ . A combined collection of 8,008 mouse genes, detected as differentially expressed in contrasts involving control and mitochondrially dysfunctional cell lines, was assembled to define gene clusters with the kmeans method. A second collection of 9,191 differentially expressed genes was assembled by including those from contrasts involving cells in which alternative oxidase (AOX), or ND1L (NA), had been introduced to rescue a mitochondrial defect; this collection was used to compare the transcriptomic profile of all mouse cell lines with PCA. Similarly, a combined collection of 8,165 differentially expressed human genes was selected for the construction of k-means clusters and, also, to compare transcriptomic profiles with PCA

**Table S2.**
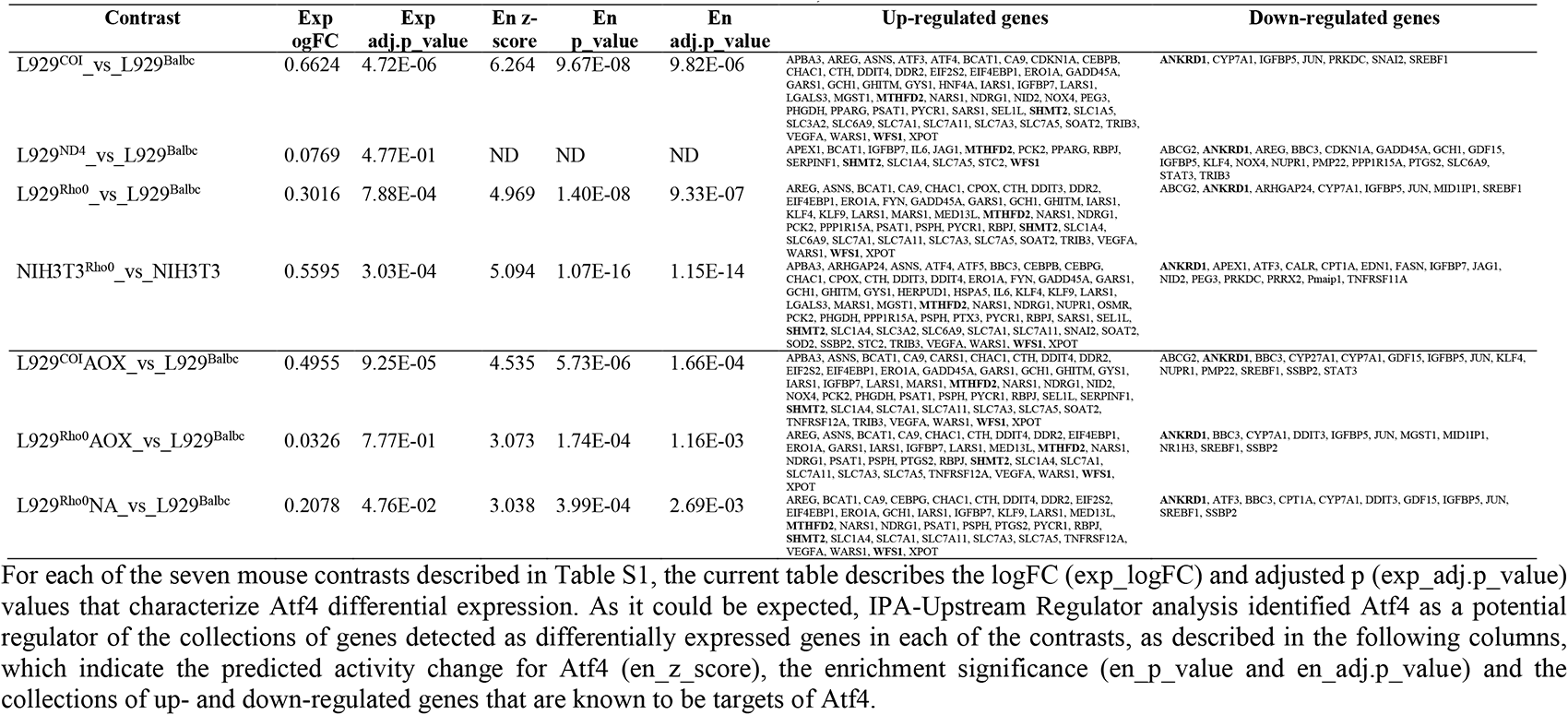
Summary Atf4-related IPA-Upstream Regulator analysis results, in mouse.

