## Supplementary figures and images for "An ATF4-centric regulatory network is required for the assembly and function of the OXPHOS system"

### Supplemental figure 1

#### K-mean 30 Mouse Clusters

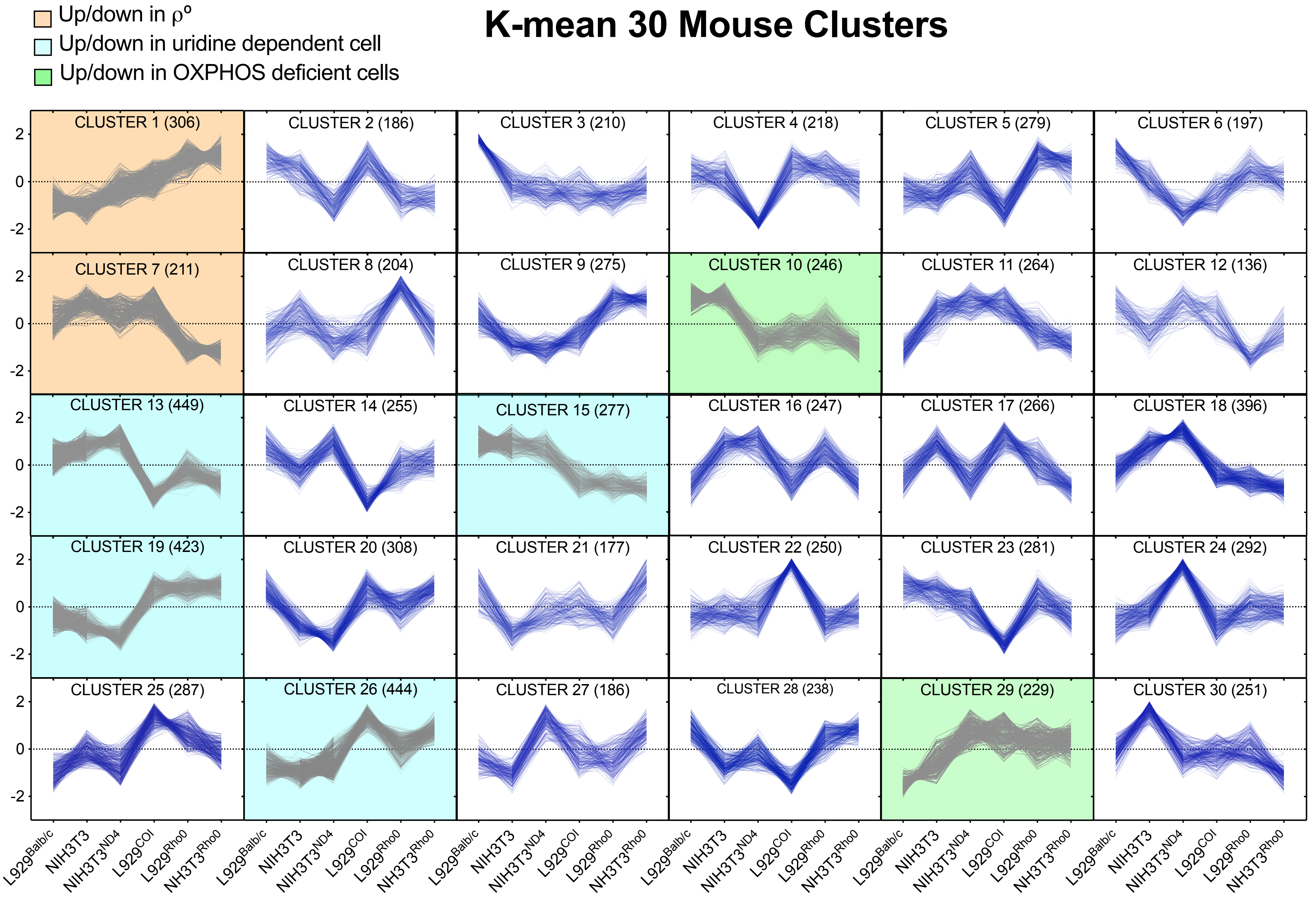

### Supplemental figure 2

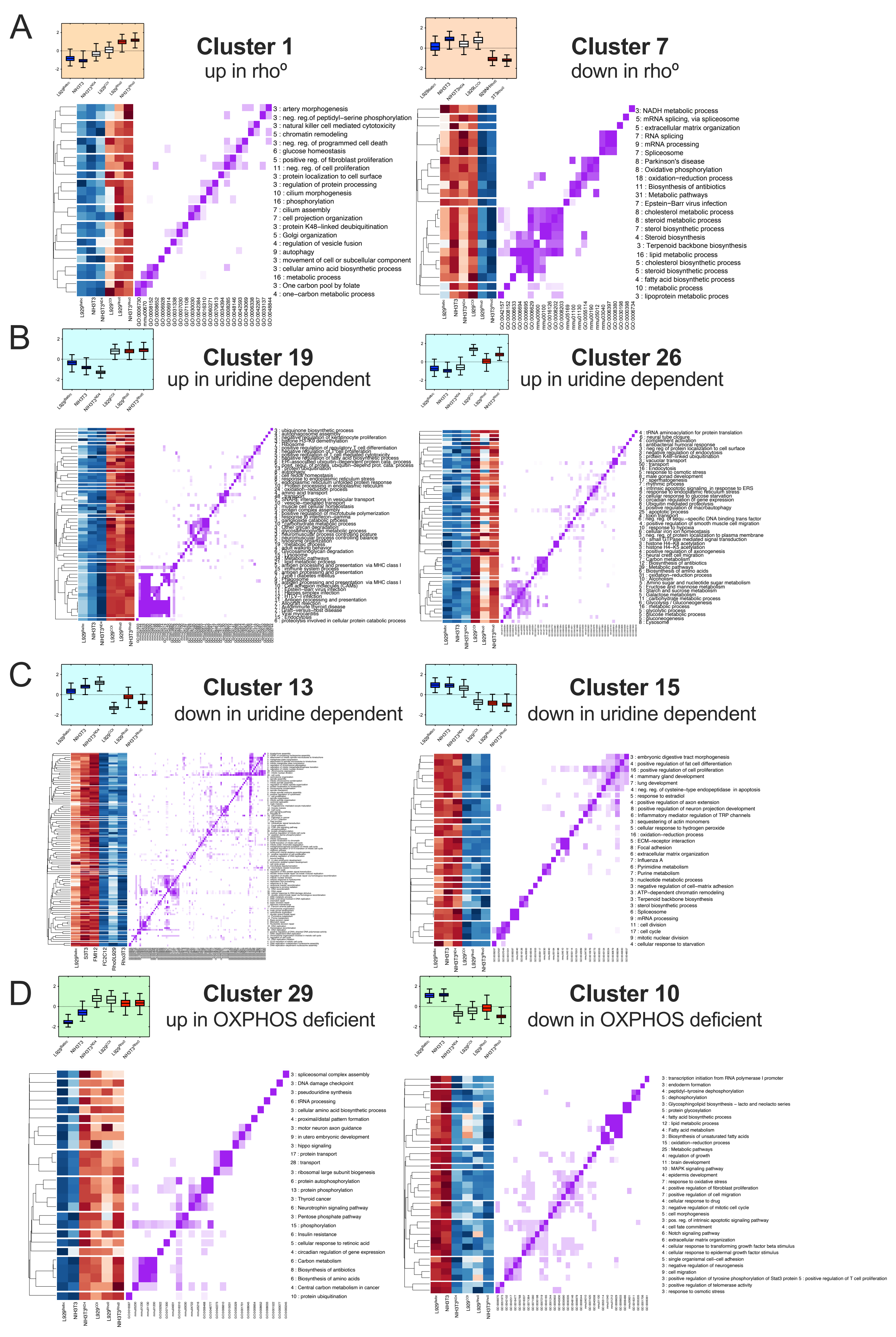

### Supplemental figure 3

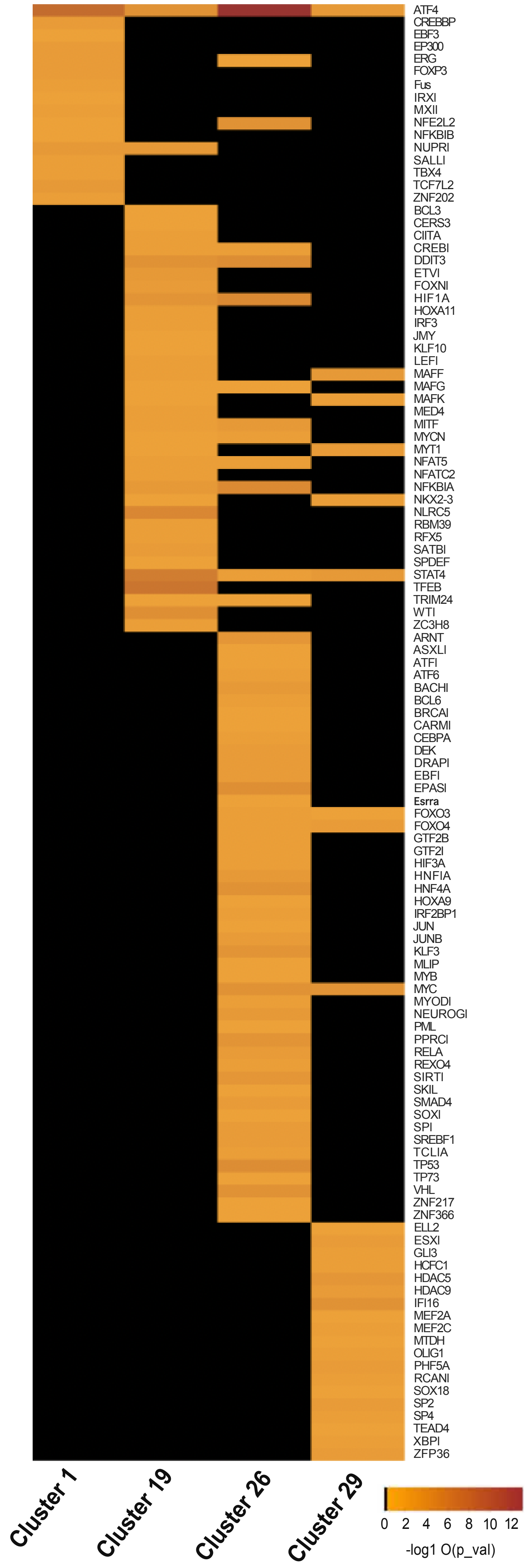

### Supplemental figure 4

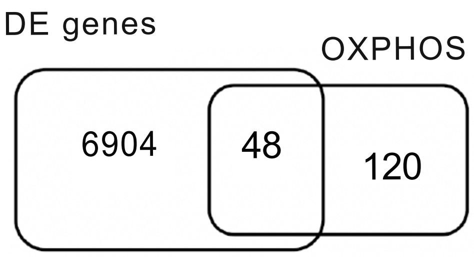

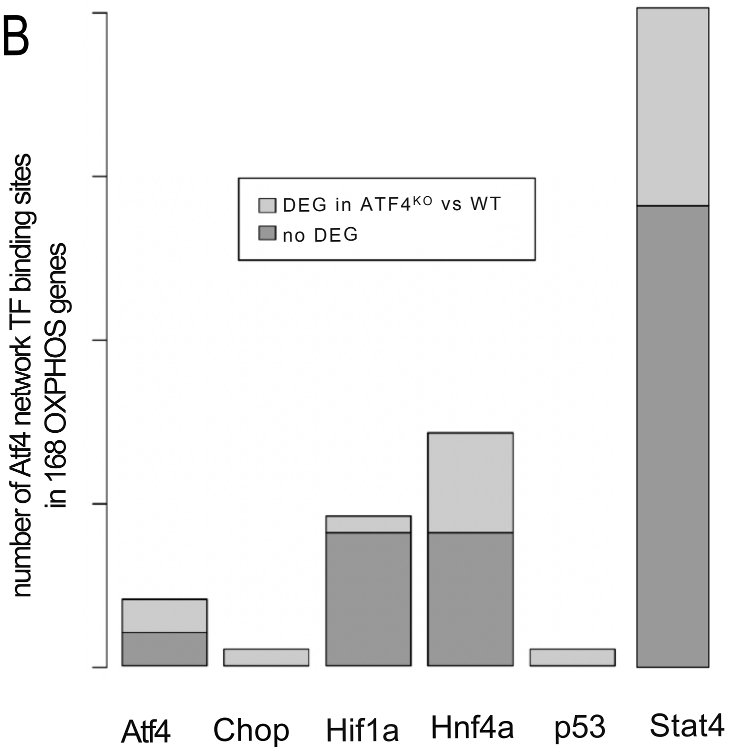

### Supplemental figure 5

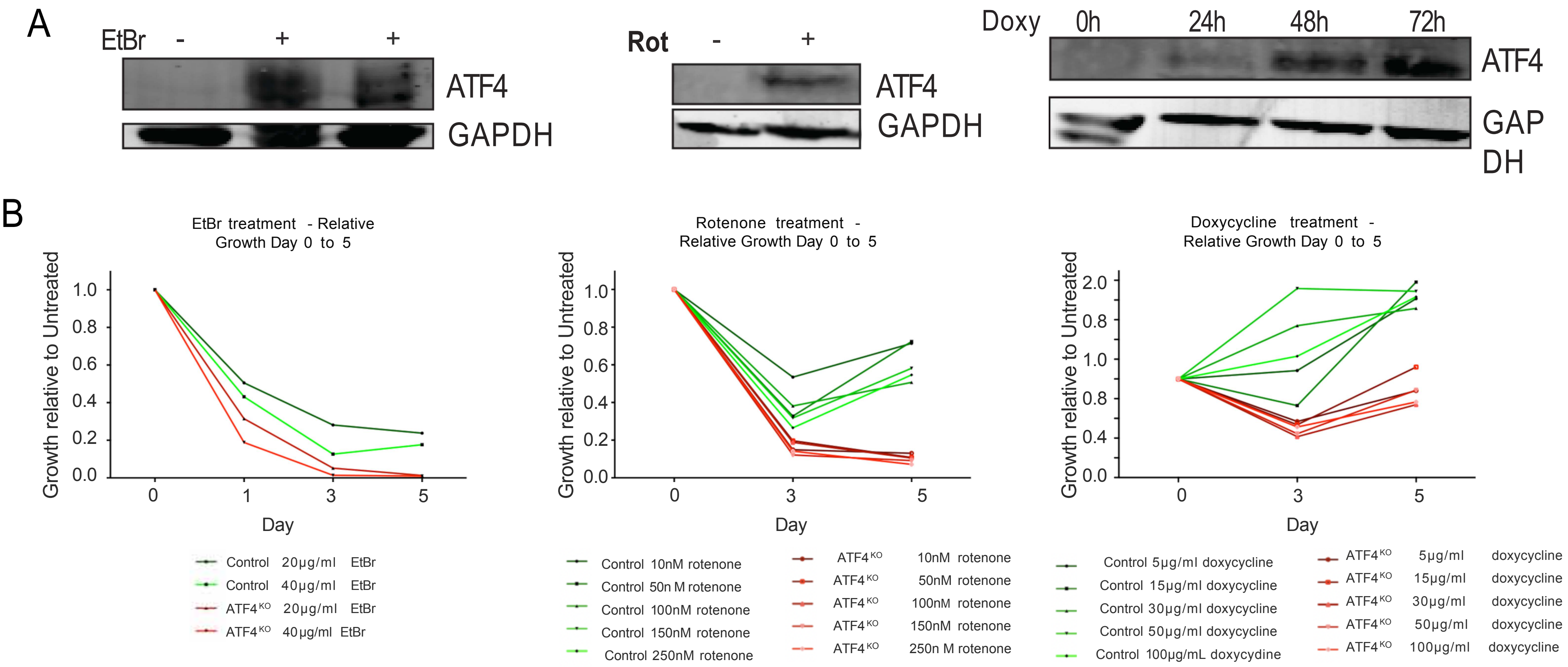

### Supplemental figure 6

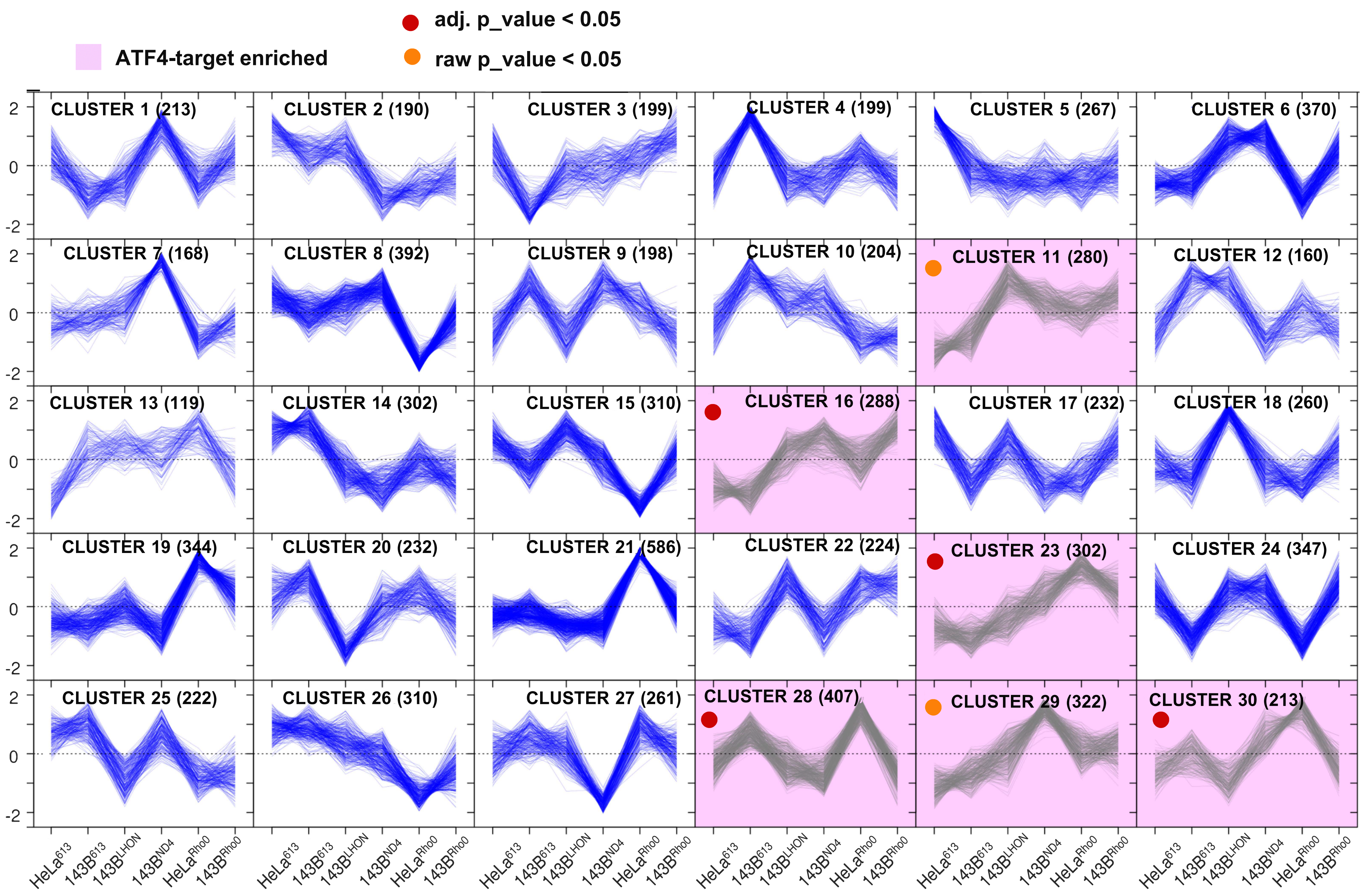

### Supplemental figure 7

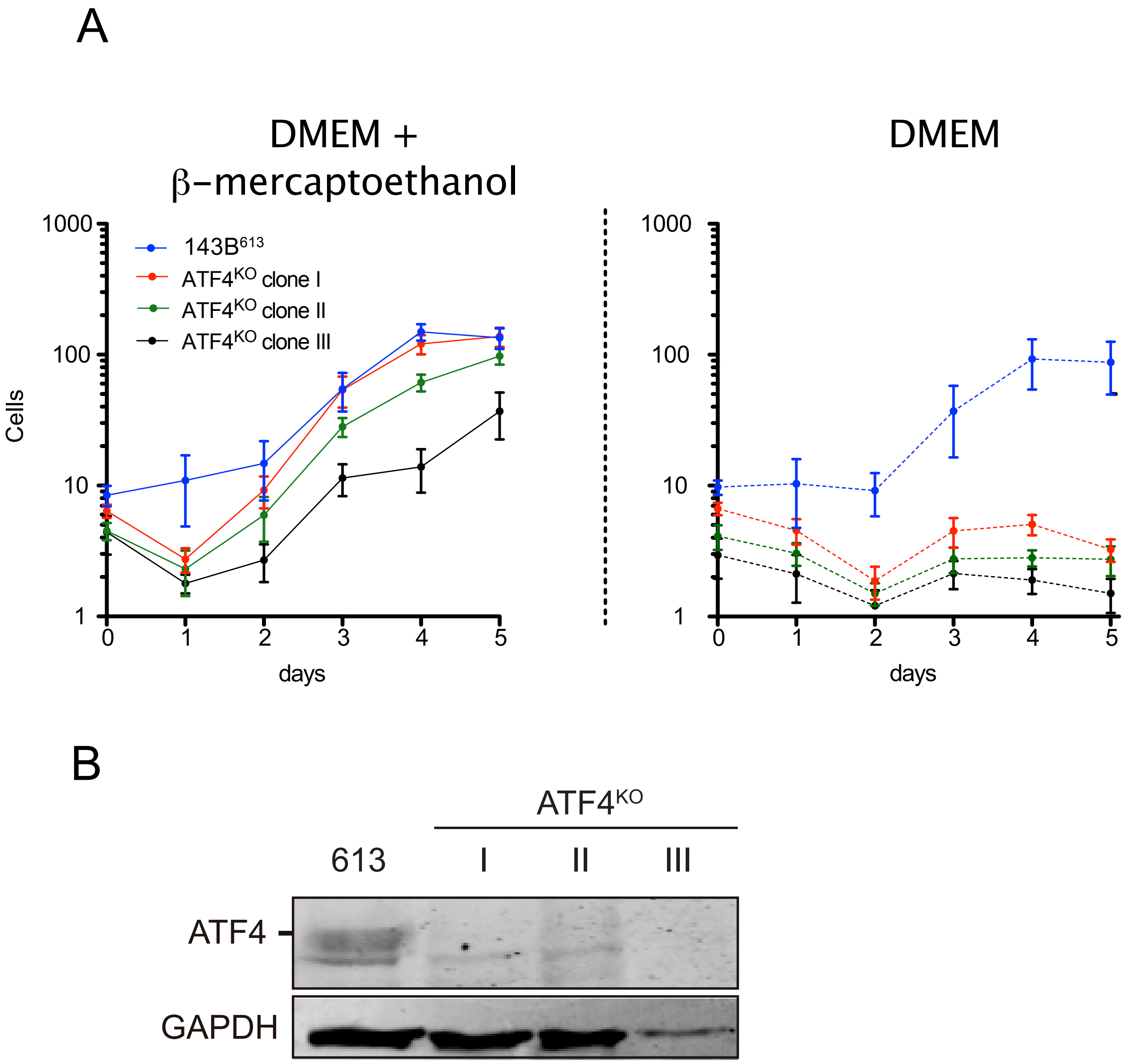


Figure S7. Validation of CRISPR/Cas9 colonies
